# Spatiotemporal variation in cutin polymerization and remodeling mediated by GDSL-hydrolase enzymes during tomato fruit development

**DOI:** 10.1101/2025.01.09.632122

**Authors:** Glenn Philippe, Iben Sørensen, Aurore Guérault, Marcella J. Cross, David S. Domozych, Mads H. Clausen, Jocelyn K. C. Rose

## Abstract

Land plants produce a cuticle, an extracellular hydrophobic layer that covers aerial organs and is involved in many critical protective roles, most notably in preventing desiccation. The predominant component of the cuticle is the lipidic polyester, cutin, which is deposited in the epidermal primary cell wall. Most of cutin of tomato fruit, a model for cuticle research, is polymerized by the extracellular GDSL-hydrolase enzyme CUTIN SYNTHASE-LIKE 1 (CUS1). However, other enzymes involved in cutin assembly remain to be identified and characterized. In this current study, we investigated whether other GDSL-hydrolases that are highly expressed in fruit epidermis might also contribute to cutin polymerization and restructuring. Candidates include homologs of *Arabidopsis thaliana* CUTICLE DESTRUCTIVE FACTOR 1 (CDEF1), which has been reported to catalyze cutin hydrolysis, as well as other phylogenetically diverse and distantly related GDSL-hydrolases. We determined that members of the CUS and CDEF families can catalyze the transesterification of cutin precursors *in vitro*, and can modify tomato fruit cutin structure in semi-*in vivo* assays. Tomato mutant knockout lines of *CUS* and *CDEF* genes generated by CRISPR/*Cas9* and cross mutations with *cus1* (previously *cd1*) were found to exhibit different fruit and flower phenotypes related to cutin assembly, including an effect on cutin monomer esterification, composition and content, cutin nanoridge formation in flowers, fruit cuticle permeability and permeance. Characterization of the mutant phenotypes, in combination with the enzyme analysis and bioassays, revealed distinct differences in the contribution of CUS and CDEF enzymes to cutin biosynthesis and remodeling. Our analysis also revealed unexpected spatiotemporal variation in cutin polymerization and structure coordinated by distinct GDSL-hydrolase enzymes over the fruit surface, which further suggests great complexity in cutin deposition and cuticle functions during organ development.

**Highlights:**

- Cutin polymerization in tomato is catalyzed by coordinating the spatiotemporal expression of CUTIN SYNTHASE enzymes in different organs, including during fruit development.
- Extracellular cutin polymerization is not a function limited to the canonical CUTIN SYNTHASE family members but can be also be catalyzed by other GDSL-hydrolase enzymes, as suggested by evidence *in vitro*.
- Tomato CDEF enzymes, a clade within the GDSL-hydrolase superfamily, are involved in remodeling cutin structure during fruit development.
- The biosynthesis and remodeling of cutin over the tomato fruit surface is spatially heterogeneous.

## Introduction

The plant cuticle, which is mainly localized at the outermost part of primary cell walls of epidermal cells, serves as a barrier protecting aerial plant organs and preventing water loss. It is composed of cutin, an insoluble polyester of glycerol, phenylpropanoids, hydroxy and epoxy fatty acids, which is entangled with polysaccharides and fulfilled by a broad range of waxes. In tomato (*Solanum lycopersicum*), the main cutin monomers are C16-derived fatty acids (9/10,16-dihydroxyhexadecanoic acids and 16-hydroxyhexadecanoic acids), which are some of the most commonly spread monomers in plants (Fich et al. 2016). Like many fleshy fruits, tomato fruits develop a thick astomatous cuticle that influences commercially important features such as susceptibility to fruit cracking, visual appearance, and post-harvest shelf-life (Martin and Rose, 2014). Therefore, it has emerged as a model for studying cuticle formation and properties. Insights into the regulation, synthesis and transport of precursors, as well as structure and properties of the cuticle have been obtained in tomato (Petit et al. 2021). However, the CUTIN SYNTHASE 1 (CUS1) represents the only characterized enzyme involved in the extracellular assembly process of the cuticle to date (Yeats et al. 2012; Girard et al. 2012). Tomato plants affected in *CUS1* gene expression presented severe reduction of cutin deposition in fruits, which originate from defect of its polymerization. The enzymatic mechanism leading CUS1 to polymerize cutin has been subject of different studies, providing important knowledge to understand the formation of the cuticle (Philippe et al. 2016; San Segundo et al. 2019; Yeats et al. 2014). CUS1 is a member of the GDSL-hydrolase superfamily and belongs to the CUS family, for which orthologs are found in evolutionary distant land plant species as far as bryophytes (Yeats et al. 2014). Function of transesterification of 2-MAG cutin specific precursors is conserved through the gene family, which lead to linear and cross-linked structural patterns of cutin (Yeats et al. 2014; Philippe et al. 2016; San Segundo et al. 2019). Such research have raised many new questions about other existing mechanisms for extracellular formation of the cuticle that involves a complex and dynamic assembly of many components, as well as remodeling mechanisms allowing cuticle integrity while expansion of plant organs that remained undetermined (Xin et al. 2021). Particularly, during fruit growth the cuticle must remain both flexible and intact but also retain its barrier properties.

Theories have emerged including questionable and unproven cutin self-assembly (Segado et al. 2020), however the existence of many uncharacterized enzymes and the possibility to confirm their effect through gene mutations or to test them through *in vitro* assays determines a trustworthy research area in enzymatical biosynthesis. Candidate genes include the cryptic *BODYGUARD* for which function remains unclarified years after a cuticular phenotype was observed in Arabidopsis, as well as members of the GDSL-hydrolases gene superfamily that forms an interesting cluster of cuticle-related enzymes. Although the wide diversity of GDSL-hydrolases displays multifunctionality involved in essential roles in all kingdoms, including regulation of plant growth, development, and responses to abiotic and biotic stresses (Shen et al. 2022; Akoh et al. 2004), a few members not belonging to the CUS family have been described for their activity associated with the cuticle in Arabidopsis. The *Cuticle Destructing Factor 1* (*CDEF1*), for which ectopic overexpression causes cutin degradation, and the *Occlusion of Stomatal Pore 1* (*OSP1*) that is associated with stomatal cuticular ledge formation, display hydrolytic activities (Takahashi et al. 2010; Tang et al. 2020). Several enzymes were also identified to be related to the chemically close polyester to cutin, suberin, for which redundant polymerization and hydrolysis functions were described (Ursache et al. 2021). In the distant Poaceae species barley and rice, mutants of *HvGDSL1* and *OsWDL1* exhibit reduced or ultrastructural defects of the leaf cuticle causing important water loss, though a function in the deposition of cutin remains speculative (Li et al. 2017b; Park et al. 2010).

Several studies performed phylogenetic and gene expression analyses of the GDSL-hydrolase superfamily in Arabidopsis as well as in various cultivated plants such are herb, cereal and berry plant, creating supported theoretical basis for the functional prediction of many genes and selection of candidate genes for further detailed functional study in these species (Chepyshko et al. 2012; Lai et al. 2017; Li et al. 2019; Ni et al. 2020; Huo et al. 2020). In the present study we performed the tomato GDSL-hydrolase phylogenetic tree in order to highlight putative relationships with the previously described genes, then we established a selection of candidates based on fruit epidermis gene expression collected in the database of the tomato expression atlas project (Shinozaki et al. 2018). Recombinant enzymes of highly and region-specific expressed genes were produced to investigate functional relationship with cutin precursor and native cutin through *in vitro* and newly developed semi-*in vivo* bioassays. Members of the CUS and the CDEF family were then chosen to be studied through the generation of CRISPR/*Cas9* lines and breeding with *cd1* (renamed *cus1*) genomic background for deep phenotype characterizations. in already affected. With the combination of diverse approaches, we identify a cutin remodeling enzyme and show a spatio-temporal coordination of the cutin biosynthesis by GDSL-hydrolases in tomato.

## Materials and Methods

### 1. Gene candidates screening

#### 1.1. Quantitative PCR

Quantitative PCR (qPCR) analysis of the tomato CDEF genes was performed on outer epidermis of 4 freshly harvested fruits at 15 dpa (days post-anthesis). 2 disks of 1 cm diameter were collected on fruit surface with a cork borer on the 3 targeted regions. Expression was normalized to *EF* gene.

#### 1.2. Phylogenic tree

The last version of the tomato genome Itag04.0 (https://solgenomics.net/) was used to blast CUS1 amino acid sequence with BioEdit Software to gather GDSL proteins, then the candidates were tested against the Hidden Markov Model (HMM) database (https://www.ebi.ac.uk/Tools/hmmer/search/hmmscan) for GDSL domain profiles numbered PF00657 and PF13472. Subsequently, 5 genes were excluded from our analyses. Characterized GDSL enzyme sequences from other organisms were then added to the pool before alignment with M-coffee method (https://tcoffee.crg.eu/) and building of q Maximum Likelihood tree with 100 bootstraps and standard parameters by MEGA11 software. Non tomato enzyme sequences include *Arabidopsis thaliana* CUTIN SYNTHASE 1 and 2 (*At*CUS1 and *At*CUS2), CUTICLE DESTRUCTING FACTOR 1 (*At*CDEF1), OCCLUSION OF STOMATAL PORE 1 (*At*OSP1), EXTRACELLULAR LIPASE 4 (*At*EXL4), GDSL LIPASE 2 (*At*GLIP2), ALPHA-FUCOSIDASE 1 (*At*FXG1), 10 suberin-related genes renamed SUBERIN SYNTHASE, SUBERIN HYDROLASE or SUBERIN RELATED for clarity according to their described functions (Ursache et al. 2021) (*At*GELP96, *At*GELP38, *At*GELP51, *At*GELP49 and *At*GELP22 renamed *At*SUS1, *At*SUS2, *At*SUS3, *At*SUS4 and *At*SUS5 respectfully; *At*GELP55, *At*GELP12, *At*GELP72 renamed *At*SUH1, *At*SUH2, *At*SUH3 respectfully; *At*GELP73 and *At*GELP81 renamed *At*SUR1 and *At*SUR2 respectfully), as well as *Ipomoea batatas* ISOCHLOROGENATE SYNTHASE (*Ib*lCS), *Medicago sativa* EARLY NODULINS 8 (*Ms*ENOD8), Rice WILTED DWARF AND LETHAL 1 (*Os*WDL1), Barley GDSL1 (*Hv*GDSL1), *Digitalis lanata* LANATOSIDE 15-O-ACETYLESTERASE (*Dl*LAE), *Agave americana* and *Physcomitrella patens* CUTIN SYNTHASES (*Aga*SGNH and *Pp*CUS1 respectfully). Motif homology search was performed by Multiple Em for Motif Elictitation (https://meme-suite.org) and added to the tree via the design tree website itol (https://itol.embl.de).

Expression value and pattern of the selected tomato GDSL-hydrolase candidates were obtained from the Tomato expression atlas (https://tea.solgenomics.net/expression_viewer) for fruit and from the Tomato eFP Browser (http://bar.utoronto.ca/efp_tomato/cgi-bin/efpWeb.cgi) for the whole plant.

### 2. Recombinant proteins activity

#### 2.1. recombinant enzyme production

The protein expression and purification were conducted as described by Yeats et al. (2014) by cloning the coding sequences into pEAQ-HT vector, agroinfiltration of in *Nicotiana benthamiana* leaves and separation of the enzymes with Ni-IMAC resin (Thermo Scientific). Size of the expressed proteins were verified by western-blot with monoclonal anti-polyhistidine produced in mouse (Sigma Aldrich).

#### 2.2. In vitro enzymatic activity

The cutin precursor substrate 2-Monoacylglycerol (2-MAG) was synthesized according to Yeats et al. (2012). Incubations of the substrate with the purified enzymes (described 2.1) were conducted in duplicates for 12h or 24h in tube or on fruit inner epidermis (see description 2.3) according to purpose, with concentrations, volumes and conditions determined by Yeats et al. (2014). Reaction products were extracted in ethyl acetate and crystallized on two spots of a MALDI target plate with 2,5-dihydroxybenzoic acid following the previous protocol (Yeats et al. 2012). Analysis was performed by autoflex maX MALDI-TOF (Bruker) and spectra of at least 3 shots per spot were treated on flexAnalysis software by extracting the specific 2-MAG mass 385.3, hydrolytic product dihydroxyhexadecanoic acid mass 311.2, and oligomeric products mass 655.5 (n=2), 925.7 (n=3), 1195.9 (n=4), 1466.1 (n=5), 1736.4 (n=6), 2006.6 (n=7) and 2276.6 (n=8). Ion abundance were then used to calculate the relative proportion of 2-MHG in conversion products.

#### 2.3. Semi-in vivo bioassay

Fresh tomato fruit around 15-20dpa were cut and opened, then locular tissue and seeds were carefully removed with no stretching or contact with the fragile inner epidermis. A light stream of distilled water was run on the fruit inner epidermis for 15 sec, then 100uL of a mixture described above was dropped and a picture was taken to further outline of the reaction. Samples were gently transferred in 100%Relative Humidity chamber for 12h incubation. The mixture was then back and forth pipetted to recover the substrate not bound to the cutin without touching the epidermis and prepared for analysis by MALDI-TOF as above. The inner epidermis was washed with a light stream of distilled water for 30sec and filled with toluidine blue (TB) 0.02% for about 15 min or until a uniform but faint blue background was obtained after a final wash. For each fruit, a locule was scarified with TB staining to control the integrity of the fragile inner epidermis and validate the use of the opposite locule for enzymatic incubation.

### 3. Plant material

#### 3.1. Plant growth

Tomato plants (*Solanum lycopersicum*) cultivar M82 (wild-type) and mutants (*cd1* renamed *cus1*, CRISPR/*Cas9* edited lines and crosses between *cus1* and the CRISPR/*Cas9* lines) were grown in a greenhouse according to Isaacson et al. (2009).

#### 3.2. Generation of CRISPR/Cas9 lines

Vector construction for targeted sequence edition is build according to (Reem and Van Eck 2019) and agrobacterium-mediated transformation was performed by Plant transformation was performed by the Center for Plant Biotechnology Research of the Boyce Tompson Institute (Ithaca, USA) for at least 10 plants. Sanger sequencing were carried out on the young T0 and T1 plants for selection of *Cas9*-free genotype with correct gene knockout mutation.

#### 3.3. Generation of multiple mutant lines

The different CRISPR/*Cas9* lines were crossed with *cus1* by manual pollination and selected in the two following generations to obtain plants with homozygous mutations in each targeted gene.

#### 3.4. Cuticle isolation and dewaxing

Fruit outer pericarp was cut from the stem, equatorial and stylar regions in discs of 2 cm diameter (transpiration chamber) and 7.3 mm (biochemical analysis) with a cork borer, as well as large rectangles of around 1 cm x 6 cm (biomechanical assays) with razor blade, and incubated in an enzymatic cocktail of 1% cellulase (Sigma) and 2% pectinase (Sigma) in 50mM citrate buffer pH 4 containing 0.02% (w/v) NaN3, on a rocker with medium agitation at room temperature. The enzymatic cocktail was changed weekly over 2 months, then the cuticles were rinsed carefully and dried for further analysis. Waxes were removed before cutin biochemical analysis, biomechanical assay, and permeance measurement when needed, through 3 successive baths of at least 5 min in chloroform.

### 4. Fruit resistance to stress

#### 4.1 Fruit water loss rate and toluidine blue staining

At least 10 red ripe stage fruit were freshly harvested from each genotype, rinsed with water and dried. Fruit with no apparent surface damages were placed in trays and set aside on an undisturbed bench at room temperature. Fruit were weighed every week over a month.

Toluidine blue O (TB) staining was performed by immersing clean fruit in a 0.02% (w/v) solution for 2h, then washed with a stream of water before imaging with a camera (Nikon) mounted on a swivel arm.

#### 4.2 Desiccation chambers-cuticle permeance

Permeances of isolated cuticles were determined according to Fich et al. (2020), using custom-built aluminum transpiration chambers. While the weight of the assembled chamber and cuticle was measured over 4 days, the weight of the assembly with dewaxed cuticle samples was measured over 24h due to high permeance. Samples with abnormal water loss due to tears or bad assembly were discarded and between 11 to 15 replicates were kept for each sample.

#### 4.3 Biomechanical assay

Tomato cuticles were isolated and dewaxed as described in 3.4. Strips with a width of 4.5 mm were cut with a sharp razor blade and these were fastened to a paper tab with double sided tape below and above the sample. The tabs were immobilized in house made 3D printed adapters on an Instron model 3342, and the sides of the tabs were cut with scissors. The Instron was set to a 10 mm fixture separation and a rate of displacement of 3 mm/min until breakage. The maximum force reached before the breaking point was measured for at least 5 replicates per sample.

### 5. Microscopy

#### 5.1 light microscopy

Tissue fixation, embedding, cryosectioning and slide preparation were performed as described by Buda et al. (2009). Then, cuticle lipidic material was stained with Oil Red O dye (Alfa Aesar) in isopropyl alcohol and viewed on an AxioImager A1 microscope (Zeiss) as described by Martin et al. (2017).

#### 5.2 SEM

Flowers were harvested 1-2 days after anthesis, and anthers (cut once lengthwise) and styles were immediately removed and fixed in a phosphate buffer containing 2.5% glutaraldehyde by vacuum infiltrating for 4 hours and storing at 4°C over night. After fixation, samples were dehydrated at room temperature in an ethanol series (30%, 50%, 70%, 80%, and 90%) for 10 min each, then two times in 95% ethanol for 20 min each. The samples were then transferred to a mixture of 95% ethanol and isoamyl acetate (1:1) for 10 min and 100% isoamyl acetate for 15 min (Bhattacharya et al. 2020). After evaporation of the isoamyl acetate, samples were placed onto scanning electron microscope (SEM) specimen stubs. The stubs were placed in boxes with Drierite (VWR) until SEM observation. SEM was performed with a Phenom XL Desktop SEM (https://www.nanoscience.com/products/scanning-electron-microscopes/phenom-xl/). Samples were sputter coated for 40 seconds using a 108 Manual Sputter Coater (https://www.tedpella.com/cressington_html/Cressington-108-Manual-Sputter-Coater.aspx) with the pressure set to 0.05 mbar. Stubs were transferred to the SEM and each sample was viewed at 2-4 locations to locate different cell types and to confirm presence or absence of cuticular ridges. Zoom levels ranged from 100-200 µm for overview images and 20-80 µm for epidermal cell images.

### 6. Biochemical analyses

#### 6.1. cutin monomers labeling

Free hydroxy groups of isolated and dewaxed cuticle discs were derivatized by benzyl etherification as described by Philippe et al. (2016). Relative proportion of terminal or midchain hydroxyl group labelling was measured for the main cutin monomer 10,16-dihydroxy hexadecenoic acid after the following depolymerization method.

#### 6.2. cutin monomer quantification

Discs of isolated and dewaxed cuticle were depolymerized with a transmethylation reaction after extensive dry of the samples. One or two discs when the samples contained *CUS1* mutation were deposited in glass tubes with 200µg or 40 µg (*CUS1* mutation) ω-pentadecalactone used as an internal standard, and immersed with 60% (v/v) methanol, 15% (v/v) methyl acetate and 25% (v/v) sodium methoxide 25 wt. % in methanol. Tubes were capped tightly and brought into heat block at 60 C for 12h, with occasional vortexing. Reaction was cooled down and stopped by neutralization with glacial acetic acid, then lipids were recovered in dichloromethane after the organic phase was washed three times with 0.9% (w/v) NaCl in pure water. The phase was subsequently dried over anhydrous sodium sulfate and evaporated to dryness under a stream of N2. The dried aliquots were derivatized by adding 50 μL each of pyridine and BSTFA and heating for 30 minutes at 70°C. The derivatized monomers were dried under N2 following the derivatization, resuspended in 0.5mL or 100uL chloroform for sample with CUS1 mutation, and then analyzed by GC using a GC 6850 (Agilent) as described by Martin et al. (2017).

## Results

### Identification of GDSL-hydrolase candidates in Solanum lycopersicum

Sequence alignments and phylogenetic tree of tomato GDSL-hydrolase enzymes have been generated with the presence of annotated GDSL-hydrolases from other species to discuss enzymatic functions (fig. 1A). As reported by (Yeats et al. 2014), the CUS proteins form an ancient family that is conserved among land plants. In tomato, it consists of 5 members that show homology with evolutionary distant CUS enzymes such as in bryophytes and in monocotyledons with the suspected AgaSGNH (Reina et al. 2007). Investigation on suberin biosynthesis shed light on the putative involvement of several members of the GDSL-hydrolases in Arabidopsis, (Ursache et al. 2021). The tree forms a SUBERIN SYNTHASE (SUS) family clade that gathers 3 *At*SUS and 5 tomato enzymes. Interestingly, despite its attributed role in suberin biosynthesis, *At*SUS4 is a close homolog of *At*CDEF1 that has been characterized for its cutin and suberin hydrolytic function (Yeats et al. 2014; Takahashi et al. 2010). We delineated a clade of 9 proteins for the CDEF family with confident bootstrap values. Although the CUS, SUS and CDEF distinct families are located in the clade VI, other enzymes supposedly involved in cutin or suberin hydrolysis are scattered in CLADE III and V (*i.e. At*OSP1, *At*EXL4, *At*SUR2, *At*SUH1, *At*SUH2 and *At*SUH3). *At*OSP1 is an enzyme involved in stomatal cuticular ledge formation (Tang et al. 2020). Mutants display a continued cuticle occluding stomatal pores, which may indicate a cutin hydrolytic function like *At*CDEF1. *At*EXL4 has an esterase activity and is required for pollen hydration on flower stigma likely by an hydrolysis of the stigma cuticle (Updegraff et al. 2009). *At*SUH have a hydrolytic function on root suberin important for root emergence (Ursache et al. 2021). *At*SUR are also involved in root emergence without a clear role toward the suberin polyester, however *At*SUR1 belongs to the SUS family and *At*SUR2 is a close *At*OSP1 homolog, both supporting their lipidic polyester related function.

**Figure 1.**
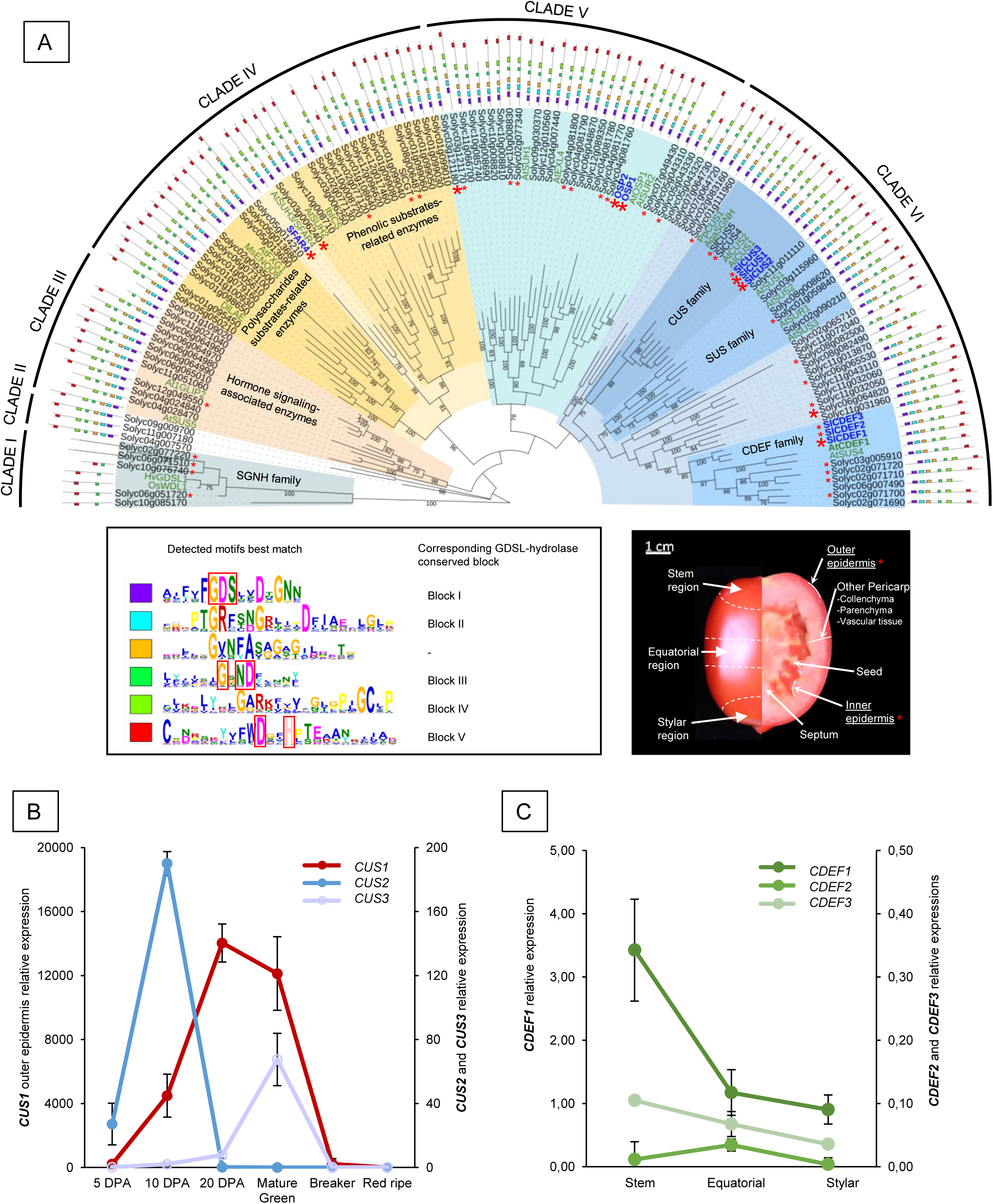
(A) Maximum Likelihood phylogenetic tree of tomato GDSL-hydrolase including a selection of GDSL-hydrolases from other species (names in green). Blue names correspond to tomato genes investigated in this study; Bold names are selected for recombinant protein production; red stars indicate genes that are expressed in tomato fruit outer and/or inner epidermis, which are indicated on the red fruit section below; big red stars indicate the highest expressed genes in tomato fruit epidermis (>100rpm value from Table 1); adjacent lines with colored squares indicates the amino acid sequence with conserved motifs indicated below and important residues are framed in red; tree branches with bootstrap value >70 are labelled. (B) Outer epidermis relative expressions of the 3 most expressed *CUS* genes throughout fruit development, values extracted from the TEA database (https://tea.solgenomics.net/). (C) Outer epidermis relative expressions of the 3 *CDEF* genes at 3 different surface areas (indicated in fig. 1A) of 15 DPA fruit, measured by qPCR.

**Table 1:**
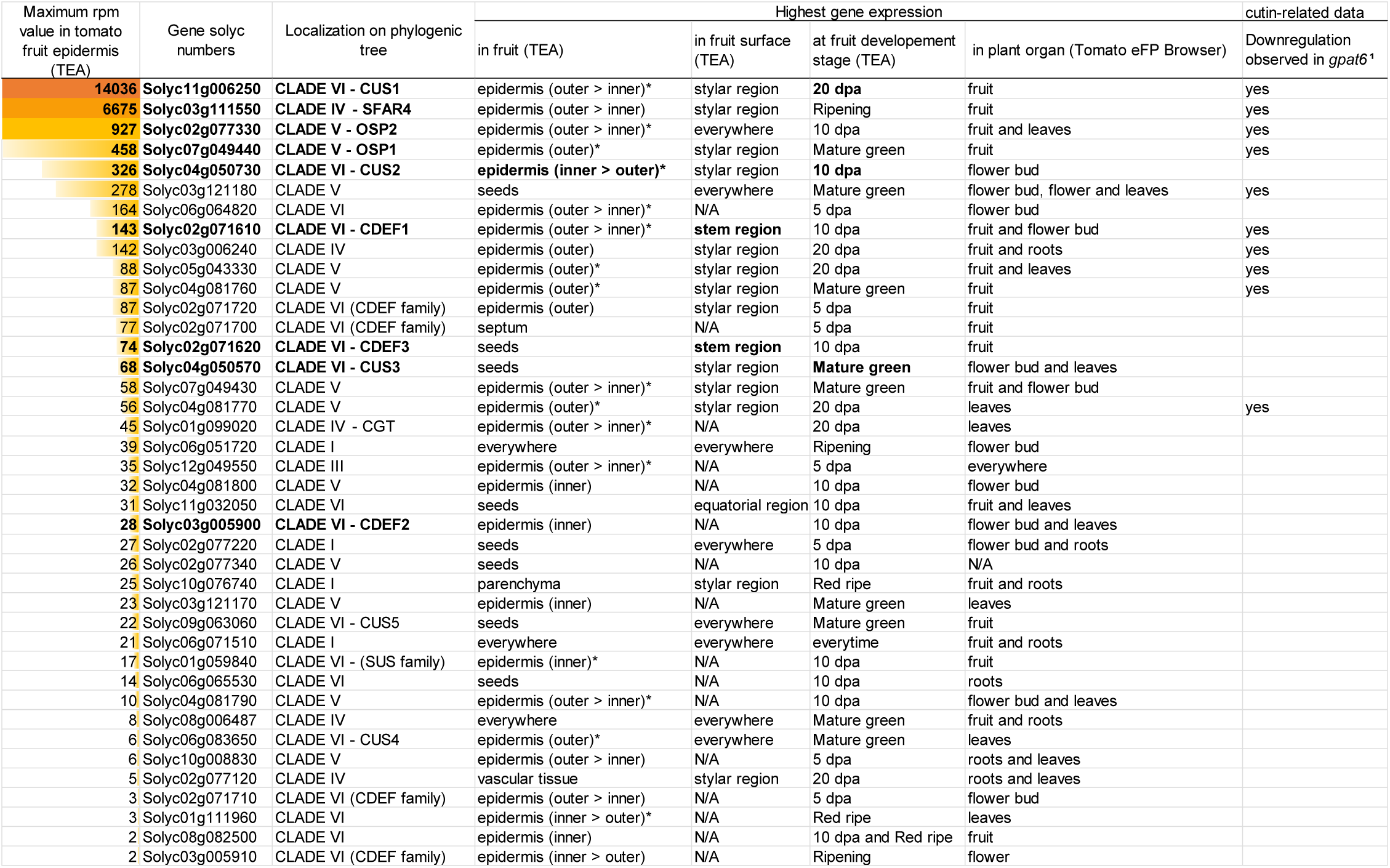

GDSL-hydrolase enzymes *Os*WDL1 and *Hv*GDSL1 have been described in rice and barley for their involvement in the cuticle integrity (Campoli et al. 2024; Park et al. 2010). However, homologs of these enzymes in tomato form the CLADE I, which regroup the SGNH-hydrolases family described by their specific GDSL motifs of the Block I and II and absence of block IV, making a distance with the other GDSL-hydrolases (Akoh et al. 2004). These enzymes, annotated isoamyl acetate-hydrolyzing esterase in the database, have no tissue specific expression in fruit and therefore are not serious candidates for cuticle-related function despite of their characterized orthologs.

On the 107 tomato GDSL-hydrolases, 40 show a gene expression in inner and/or outer fruit epidermis, which correspond to the tissues producing cutin (table 1). Although CUS1 has the highest expression by far, relatively high gene expression in tomato epidermis are observed for GDSL-hydrolases from Clade IV, V and VI. The second most expressed GDSL-hydrolase correspond to SFAR4, a tomato ortholog to *AtSFAR4/AtSUH3* that has hydrolytic function toward fatty acid (Huang et al. 2015). Surprisingly, the amino acid sequence is deprived of the GDSL block V, which is a part of the catalytic triad is a common feature of the GDSL-hydrolase superfamily members including its *At*SFAR4 ortholog (fig. 1A). The enzyme is located in the Clade IV, between two subclades of enzymes with activity related to phenylpropanoid and cell-wall-related substrates, determined by the presence of the CGT and *Ib*ICS enzymes characterized for their caffeoyl transferase activity (Miguel et al. 2020; Teutschbein et al. 2010), and *Ms*ENOD8, *At*FXG1 and *Dl*LAE identified with activity toward cell-wall substrates (De la Torre et al. 2002; Pringle and Dickstein 2004; Kandzia et al. 1998). The relatively high expression of SFAR4 in several tomato fruit tissues aligns with a substrate that is nonspecific to cutin (sup. fig. 1). The third and fourth most expressed genes are located near *At*OSP1 in the CLADE V, which regroup several cutin/suberin hydrolase enzymes previously cited. We named them OSP1 and OSP2 according to their close homology with *At*OSP1, even though this gene is involved in stomatal formation while tomato fruit is stomata-free, thus indicating a questionable role for this enzyme family. The fifth most expressed GDSL-hydrolase is *CUS2*, which differ from *CUS1* by a distinct earlier expression pattern, higher in inner than outer epidermis, and primarily located in flower development (Table 1; fig. 1B). Interestingly, unlike the most expressed GDSL-hydrolases in epidermis, CUS2 is not downregulated in tomato *gpat6* mutant, in which cutin biosynthesis is strongly affected via an impact on 2-Monoacylglycerol (2-MAG) cutin precursors (Petit et al. 2016). Within the top of the list, CDEF1 represents another particularity as its highest expression is located in the stem region of the fruit, unlike the other GDSL-hydrolases that have no or stylar region prime expression location. Moreover, diverse characterized enzymes known for their involvement in the cutin biosynthesis pathway have slight propensity to express in stylar region during tomato development such as *LACS, CYP77A, CYP86A, GPAT6, DCR* or *ABCG32*, indicating a trend for not uniformed and highly active cutin synthesis machinery at the bottom of the fruit. The peculiar region specificity was confirmed by qPCR for CDEF1, where the expression is three time more important in stem than other area, while its close homologs had not enough expression to determine a similar pattern (table 1, fig. 1C).

### Polymerization VS hydrolysis, identification of the cutin remodeling enzyme

Data obtained in a previous study indicated a polymerase activity of CUS enzymes (CUS1, *At*CUS1 and *Pp*CUS1) and hydrolytic activity of *At*CDEF1 toward the cutin precursor 2-MAG measured *in vitro* (Yeats *et al*. 2014). The epidermis high expressed GDSL-hydrolase were tested as candidates in similar *in vitro* experiments to evaluate their ability to interact with the tomato cutin precursor 2-MAG (fig. 2). We also produced closest homologs of CUS2 and CDEF1 to determine function specificity (fig. 2A). The hydrolytic activity of *At*CDEF1 toward the 2-MAG substrate is strong, with 100% conversion to 10,16-diOH-C16 acid as observed previously by Yeats et al. (2014), and absent from other enzymes with putative hydrolytic activity (fig. 2B). However, cutin polymerization products were obtained by the activity of CUS and, surprisingly, CDEF enzymes (fig. 2B). CDEF1 converted a comparable amount of substrate to oligomer products as CUS1, however with a higher proportion of dimers (fig. 2B, sup. fig. 2A). As expected, CUS2 had shown a polymerizing activity, however it remained remarkably low in comparison with CUS1, and no activity was observed with CUS3 in the same condition of time (fig. 2B). Doubling the incubation time had led to observe the achievable polymerization activity of the less active CDEF3 enzymes, while CDEF2 shows a similar activity to CDEF1 with notably high n=2 oligomer content (sup. fig. 2C). The activity of CUS2, CDEF1, *At*CDEF1 were confirmed by a glycerol release assay following transesterification or hydrolytic activity on 2-MAG substrate (sup. fig. 2D). Interestingly, the result of the enzymatic incubation performed semi-*in vivo* (*i.e*. directly on a native cutin support that is the fruit inner epidermis) led to differences between CUS1 and CDEF1 (fig. 2C). Although the reaction mixture containing CDEF1 and the 2-MAG substrate led to produce oligomers, which are detected after incubation, no 2-MAG nor oligomers were recovered from the reaction mixture containing initially CUS1 and 2-MAG. This result suggests that 2-MAG was fully polymerized to the native cutin by CUS1, while CDEF1 may have a lower polymerization activity with the native cutin or is also able to release oligomers from it in a dynamic remodeling activity.

**Figure 2.**
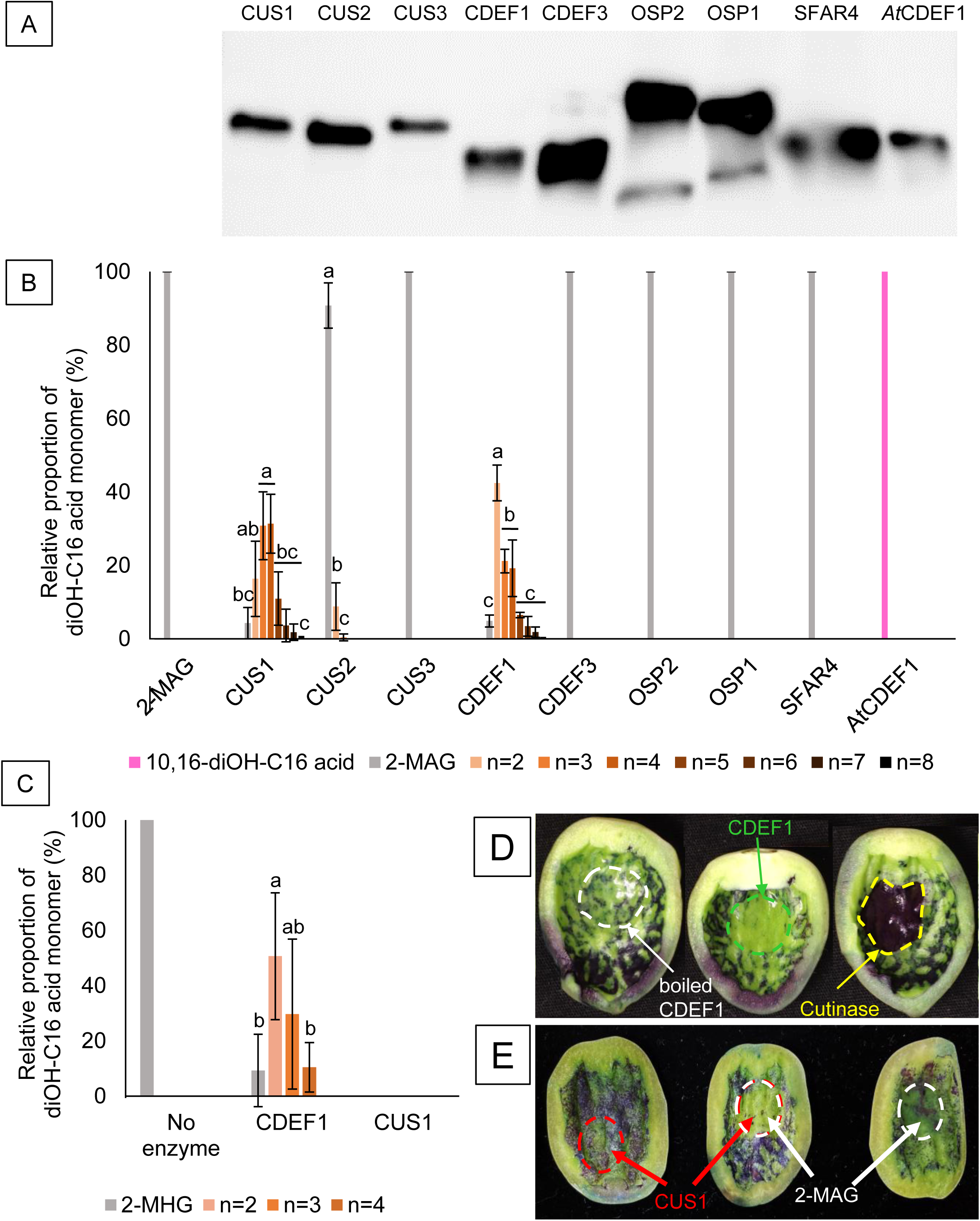
Measurement of recombinant GDSL-hydrolases activity with cutin and cutin precursors. (A) Western-blot of purified recombinant proteins. (B) 12h substrate conversion of 2-monoacylglycerol (2-MAG, light blue bar) to oligomers (orange bars) or fatty acid (10,16-diOH-C16 acid, dark blue bar) by purified GDSL-hydrolase enzymes measured by MALDI-TOF. Hydrolysis of 2-MAG is only observed with *At*CDEF1, while polymerization products are observed with CUS1, CUS2 and CDEF1 in different proportion (quantified until n=5). (C) Incubation of 2-MAG directly on native cutin from 15dpa tomato fruit. No product analyzed from the reaction mixture with CUS1 indicates a full substrate polymerization to cutin substrate *in situ*, while CDEF1 reaction mixture contains *ex-situ* polymerization product. (D and E) Bioassay for cutin remodeling activity screening. Inner epidermis of cut-in-half 15dpa tomato fruit are 12h incubated with purified recombinant enzymes CUS1 and CDEF1 (dotted frame in red), commercially pure *Fusarium solani* cutinase (dotted frame in blue), boiled CDEF1 for control (dotted frame in yellow), and 2-MAG substrate (dotted frame in white), then stained with toluidine Blue. Letters denote significant differences in enzymatic products (p<0.05) calculated by ANOVA with Tukey HSD test separated for each enzyme.

To take this analysis one step further, we stained the native cutin after incubation with enzymes. The inner epidermis of young tomato fruit has an extremely thin cutin layer, which is permeable to toluidine blue in discontinued areas (fig. 2D, E). As anticipated, incubation of a purified fungal cutinase on the substrate had led to increase the permeability of the inner epidermis by hydrolyzing the cutin, which is observed by a strong toluidine blue staining (fig. 2D). However, CDEF1 has replenished the inner epidermis with a continuous layer by actively restructuring the cutin polymer, supporting a remodeling activity. Incubations of CUS1 on native cutin with 2-MAG were also consolidated the inner epidermis, validating the role of cutin in patching holes of permeability (fig. 2E). Moreover, despite CUS2 has a lower activity toward the 2-MAG substrate, its activity on the native inner epidermis seems as strong or higher than CUS1, which is not surprising considering its specific expression pattern at this location (table 1, fig. 2B, sup. fig. 3). Interestingly, the native cutin had remained unaffected after incubation with *At*CDEF1 despite its characterized cutinase-like function (sup. fig. 3). This result corroborates the preferential hydrolytic activity toward single 2-MAG than polymerize products (sup. fig. 2B), which is in contrast to true cutinases that hydrolyse cutin oligomers (Bhunia et al. 2018).

### Generation of CRISPR/Cas9 lines and selection of cus1 double mutants to increase cutin-related phenotypes

Following the enzymatic activity observed *in vitro*, two distinct genotype lines edited by CRISPR were selected for both *CUS2* and *CDEF1,* leading to predicted gene knockouts (sup. fig. 4). *CDEF1* shared a perfect sequence homology surrounding the GDSL site with *CDEF2*, allowing the generation of *cdef1/2* double mutants via the CRISPR target.

*CUS1* impacted plants have been studied several times using different screening methods and genetical background (Girard et al. 2012; Isaacson et al. 2009; Petit et al. 2014). The resulting phenotypes were every time comparable, the fruit exhibiting a normal development excepted a massive reduction and less polymerized cutin deposition as well as a brighter appearance. Considering the single effect on the fruit cuticle formation and no visible phenotype in other organs, the previously studied *cus1* was integrated as genotype backgrounds for selection of multiple mutation lines (Isaacson et al. 2009). No effect in fruit development or morphology was observed in all *cus2* and *cdef* lines and integration of theses mutations in *cus1* plants does not affect the classical brighter phenotype exhibited by the fruit, which is conserved in all multiple mutation lines (fig. 3).

**Figure 3.**
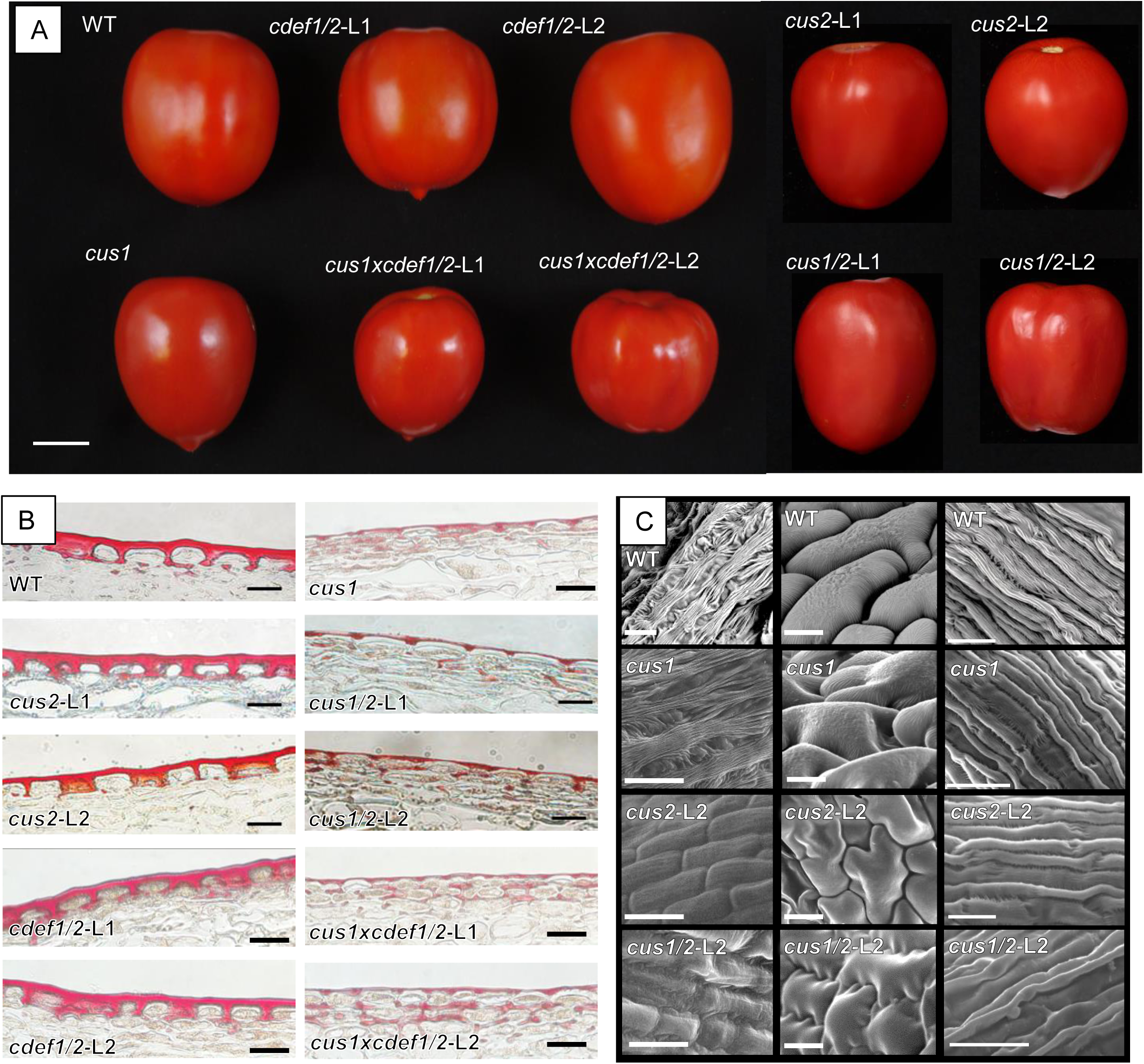
Macro and microscopic observations of generated CRISPR/Cas9 lines. (A) Red ripe stage harvested fruit; bar=2cm. (B) Fruit outer epidermis cross-sections and oil Red O staining of red ripe stage fruit for cuticle observation; bars=50µm. (C) SEM images of epidermal cells surface from flower; left column, rectangular basic cells from the upper half of anther adaxial side; middle column, basic cells from the lower half of anther adaxial side; right column, cells from the style stalk; bars=25µm.

Despite of the relatively high *CUS2* expression in fruit epidermis, cutin polymerization and composition was not effectively affected in fruit of single lines (table 1, fig. 4A). However, a lack of nanoridges was observed at the surface of flower anther cells in all *cus2* lines but not in *cus1* (fig. 3C). This phenotype has been described in characterized mutants for cutin biosynthesis in Arabidopsis (*Atcyp77a6* and *Atgpat6*) and thus remains consistent with the specifically high expression of *CUS2* in flower during their development (table 1) (Li-Beisson et al. 2009). Interestingly, nanoridges on flower style cells are completely disappearing only in the double mutant *cus1/2*, indicating putative gene compensation between *CUS1* and *CUS2* for certain type of flower cells. In fruit, the *cus1/2* lines also appeared to be valuable with an amplification of the *cus1* phenotypes described in previous studies. Namely, the overall cutin reduction was exacerbated by the double mutation with *cus2*, which was certainly resulting from the increased polymerization defect that was measured by chemical labelling of free hydroxyl groups of the main cutin monomer (fig. 4B, 5B). Furthermore, this greater phenotype results in higher cuticle permeability, which is observed by a stronger toluidine blue staining and postharvest water loss (fig. 6A-B). Surprisingly, *cus2* and *cus1/2* were not exhibited reduction of cutin content or major composition differences in the inner epidermis, where *CUS2* has its highest expression in fruit tissues (sup. fig. 5 A-B).

**Figure 4.**
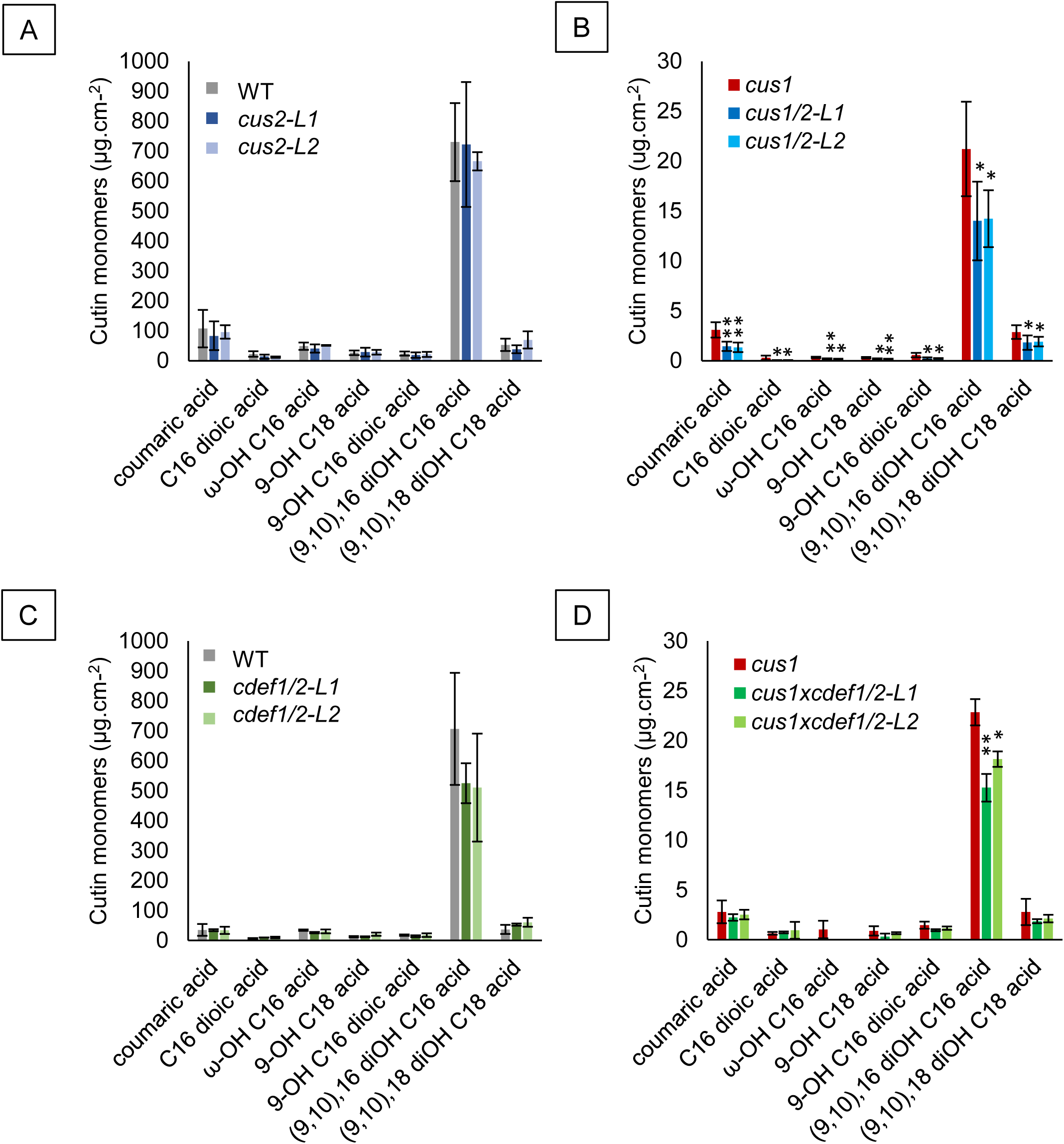
Cutin monomer composition of isolated cuticle from equatorial region (A-B) and stem region (C-D) of red ripe stage fruit. Statistical differences are indicated with asterisks (Student’s t test; *, P< 0.05; **, P<0.01; ***, P<0.001).

**Figure 5.**
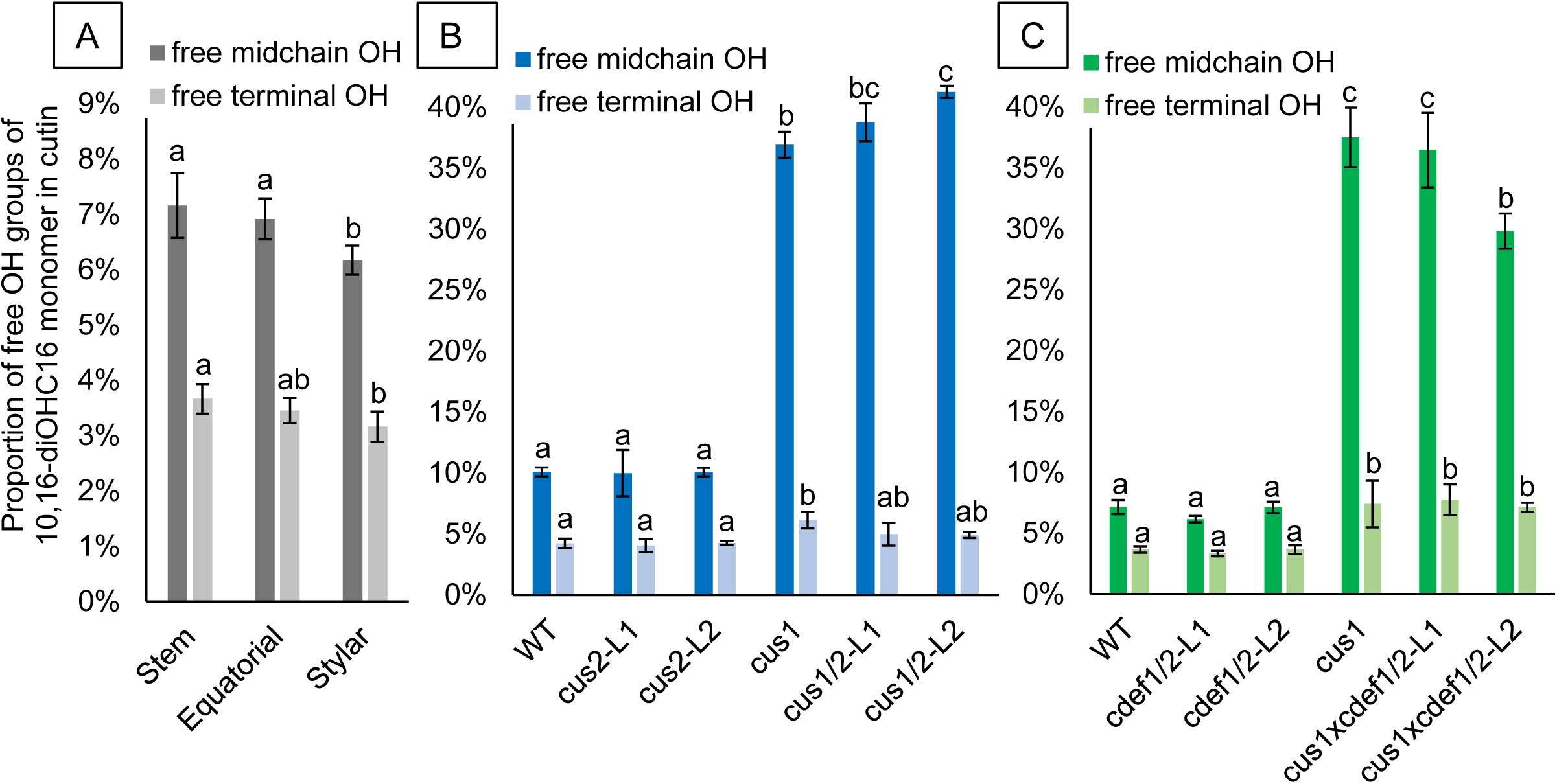
Relative proportion of non-polymerized hydroxyl groups (OH) of the major cutin monomer dihydroxyhexadecanoic acid (10,16-diOHC16) of fruit cuticle. (A) Isolated cuticle samples from different regions of red ripe stage tomato fruit. (B) Isolated cutin from equatorial area of 20dpa fruit. (C) Isolated cutin from stem region of red ripe stage fruit. Statistics are calculated by Tukey’s HSD tests following significant ANOVA results for midchain and terminal OH groups separately.

**Figure 6.**
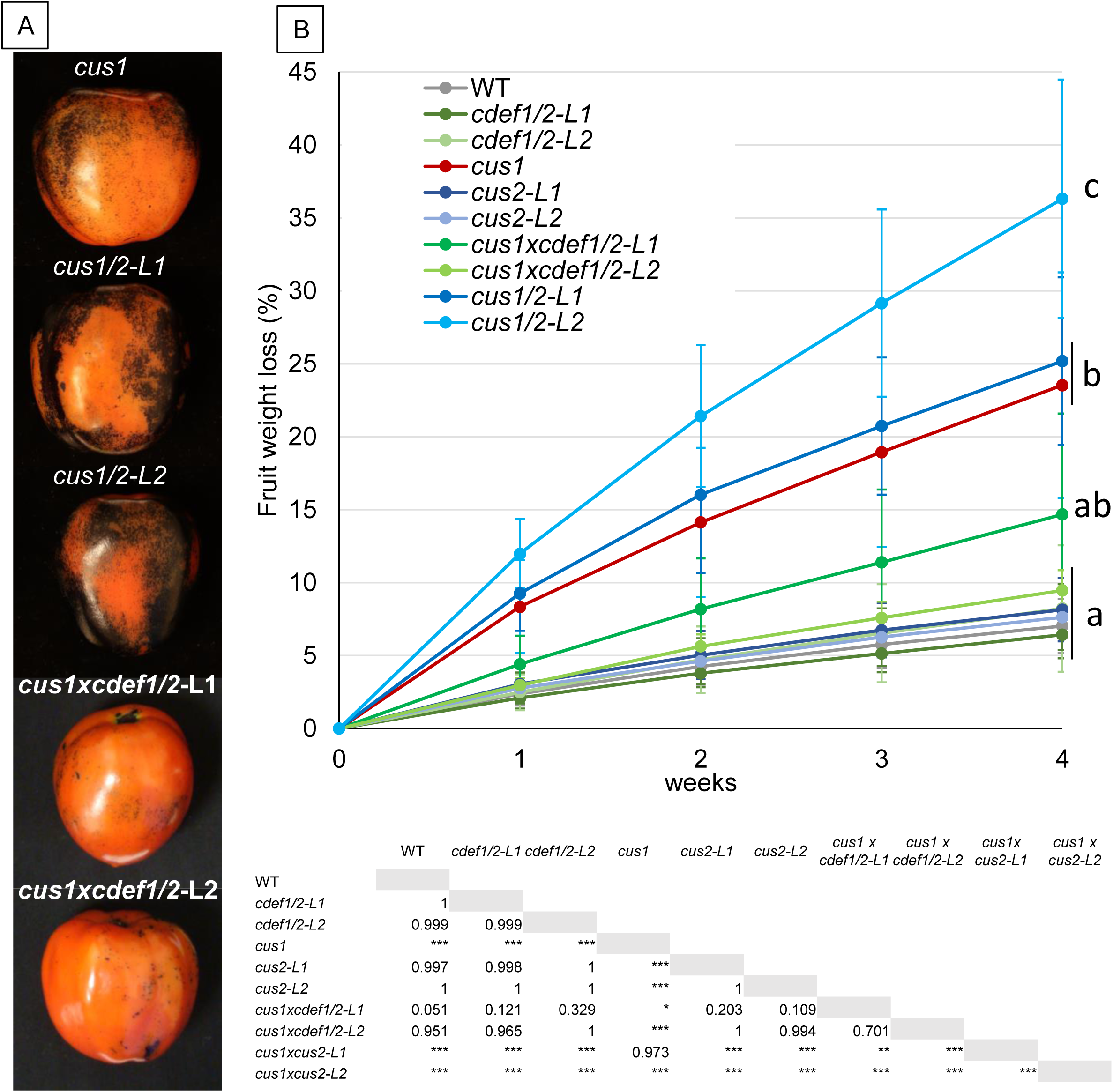
Water barrier tests of tomato fruit cuticle at red ripe stage. (A) Toluidine Blue staining of fruit reveals surface permeability. (B) Fruit transpiration measurements of freshly harvested fruit over 4 weeks. Statistical differences are calculated at the last timepoint measurement with Tukey’s HSD test; significant p-values are reported in the table as followed: *, P< 0.05; **, P<0.01; ***, P<0.001; and indicated at the extremity of the curves by letters according to a *p*-value threshold of P<0.001.

Interestingly, the degree of cutin polymerization is not homogenous along the fruit y-axis of WT tomato, for which the *CUS* genes are directly involved (fig. 5A). The proportion of non-polymerized hydroxyl groups of cutin monomers increases from stylar to stem regions, which might be a consequence of the differential expression of *CUS1* and *CUS2* that imply higher activities at the stylar region. However, it appeared that cuticle thickness and cutin content are relatively homogenous from stylar to stem regions (sup. fig. 6A-B).

An interesting feature related to the spatial cutin properties is the significantly higher permeance of dewaxed cuticle at the stem region in comparison with equatorial or stylar regions in WT (fig. 7). The permeance experiment on cuticle samples with wax did not provide significant differences, which were accounted to the important role of waxes in limiting water transpiration (sup. fig. 6C). The cuticle samples from the stem region of the WT were observed overall more fragile in this experiment with a quantity of wasted samples due to tears, which may indicate a spatial cuticle fragility (*e.g.* non-visible microcrakings). However, a decrease of cutin permeance was observed at the stem region of *cdef1/2* lines, leading to a relatively homogenous level of permeance along the fruit y-axis in the mutants. In addition, the stem region of the WT broke well in the middle sample during biomechanical assays in comparison with the *cdef1/2* lines and other region of the fruit surface, which often broke at the clamps. These observations support an innate fragility at the stem region that is modified by *CDEF1/2* knockout, although no significant differences were obtained in the data (sup. fig. 7).

**Figure 7.**
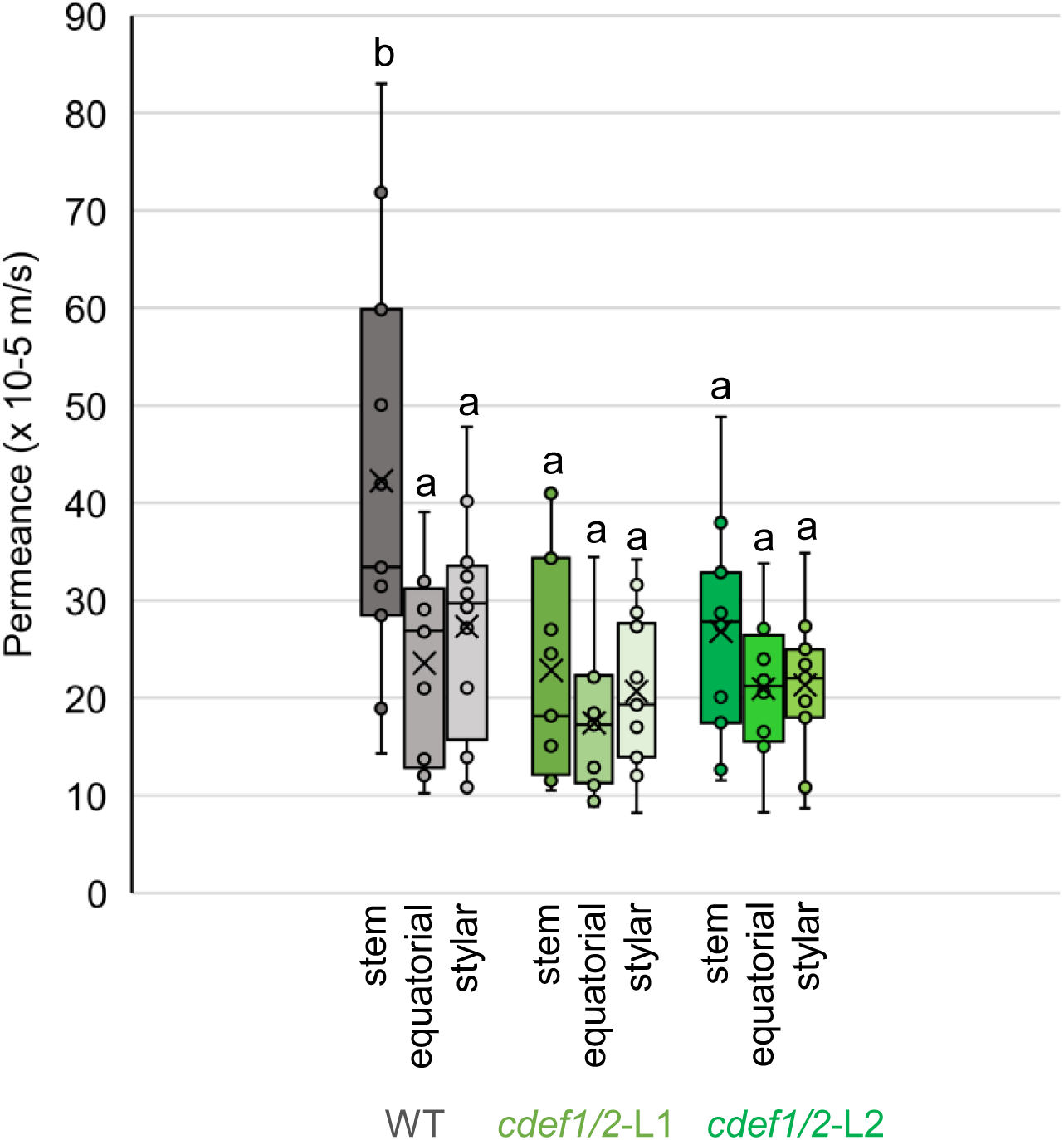
Water permeance of isolated pieces of cuticles isolated from stem, equatorial and stylar regions and dewaxed of red ripe stage fruit. Letters denote significant differences (P<0.05) calculated by ANOVA with Tukey HSD test.

Besides, the triple mutant *cus1xcdef1/2* were less affected than *cus1* in permeability, indicated by a better water retention and lower toluidine blue staining nearly rescuing the *cus1* phenotype (fig. 6A-B). Observations of cuticle cross-section revealed important deposition of cuticular material in the subepidermal cell wall layers compared with *cus1*, which may count for limiting water loss (fig. 3B). However, the analysis of the cutin composition was negatively affected, particularly in the abundance of the main monomer that decreased in *cus1xcdef1/2* stem region compared with *cus1* (fig. 4D). A slight reduction of the cutin monomer content and a thinner cuticle were also noticed in the stem region of *cdef1/2* lines, while no significant differences were observed in other regions of the fruit (fig. 4C, sup. fig. 6A-B). Measurements of cutin free hydroxyl groups in stem region of *cdef1/2* and *cus1xcdef1/2* lines had exhibited occasional differences with their respective genotype backgrounds (*i.e.* WT and cus1), suggesting occasional effect of the *CDEF* genes knockout on cutin polymerization (fig. 5C). The decrease of free hydroxyl groups indicates more cross-linking in *cdef* mutants, which is the opposite phenotype observed for *cus1* mutants. Overall, the characterization of the CRISPR lines brought evidences to a specific function of CDEF1 to the cutin assembly, particularly at the fruit stem region where the gene is mainly expressed, however radically different than CUS contribution to cutin polymerization.

## Discussion

### Distinct spatiotemporal expressions of GDSL-hydrolase genes are established for the cuticle extracellular formation of tomato

Historically, tomato had been considered as an interesting model for research on plant cuticle biosynthesis. Fruits develop smooth surface (no lenticels nor stomata) and produce a large amount of cutin with a relatively simple composition of monomers, leading to thick cuticle that is convenient to study effects of mutations in the assembly, which may lead to impressive and various cuticle phenotypes (Petit et al. 2021). However, heterogeneous chemical features have been described recently at a microscale during fruit development, revealing a complex dynamic assembly of different component of the cuticle (Reynoud et al. 2022). Our research highlighted that the three most expressed *CUTIN SYNTHASES* in fruit epidermis have successive peaks of expression during fruit development, suggesting specific steps in the cutin polymerization process (*i.e.* first *CUS2*, then *CUS1* and finally *CUS3*). The severe decrease of cutin content in *cus1* that worsens in the double mutant *cus1/2* confirms the importance of each *CUS* in the fruit cuticle formation. Furthermore, the severity of phenotype and the number of cell types affected in *cus2* flowers, also increasing in *cus1/2*, implies an effective coordinated process in different tissues and organs. These results concur with the suggested subfunctionalization of *AtCUS1 and AtCUS2* in Arabidopsis sepals from their distinct temporal expression and involvement in phenotypes (Hong et al. 2017).

We also highlighted that cutin polymerization has a naturally distinct spatial distribution on fruit surface. Cuticle heterogeneity of the whole fruit had been investigated through the study of transcuticular pores associated with dislodged trichomes, which involve a self-sealing process via localized cuticle biosynthesis or remodeling (Fich et al. 2020). Thus, the assembly requires combination of distinct mechanisms distributed at the fruit scale. The investigation of *CDEF* genes, which exhibit uncommon region-specific expression pattern, represents a good example of this complex combination of mechanisms. In this specific case, the cuticle differences at the stem region lead to question the physiological needs at this area, such a response to extended UV exposure as well as the main barrier area to transpiration. Nevertheless, the synthesis of cuticle is tightly coordinated with developmental processes, such as the formation of epidermal cells during plant and fruit growth (Javelle et al. 2011; Lashbrooke et al. 2015). Legland et al. (2012) have generated a cartography of cell elongation within mature tomato pericarp that revealed distinct cell orientations specifically associated to the stem region in the epidermis and collenchyma. The tangential direction, instead of a radial or no specific direction, indicates an internal stress and putative larger cell extension associated with fruit growth, which may trigger specific mechanisms for keeping cuticle integrity. The remodeling activity is a rational mechanism to loosen the cutin structure to this end, reminding cell wall enzymes essential for plant growth (Cosgrove 2018).

### Role of CDEF in fruit development

The complementary approaches of *in vitro* enzyme activity and characterization of CRISPR lines gears toward a complex CDEF1 function in cutin assembly that is not simply hydrolysis or polymerization. The cutin polymerization affected occasionally in a positive way in *cdef* CRISPR lines, which indicates ester hydrolysis, in addition to the esterification activity observed *in vitro,* suggests a dynamic balance controlled by *CDEF1* and leading to cutin remodeling. A comparable cutin:cutin endo-transacylase (CCT) activity has already been reported in plant epidermis extracts including tomato, but the involved enzymes remained unknown (Xin et al. 2021). The introduction of a novel semi-*in vivo* experimentations on a native cutin substrate gave results that corroborates the putative remodeling activity of CDEF1 in tomato fruit.

Interestingly, according to the characterization of mutants, CDEF1 seems to promote cutin permeability in a fruit area that might be the most exposed to environmental stress and where the cutin machinery seems less important. Whether the cutin remodeling function associated to the enzyme may explain such consequence in fruit physiology, the gene mutations had no effect on fruit shape and size, nor important cuticle damage like cracking have been observed, which may ensue from an overall biological adaptation of the plant or compensation by the numerous gene of this family expressed in fruit epidermis. Surprisingly, (Yeats et al. 2010) had already identified CDEF3 as a relatively important protein of the fruit cuticle and the second GDSL-hydrolase. However, the low expression of this homolog gene, the lower enzymatic activity, as well as the generation of phenotype-free CRISPR lines (data not shown) did not support a major role for this candidate. Although cutin permeance is known to be less correlated to cutin polymerization than thickness (Philippe et al. 2016), the CDEF enzymes have a role related to cutin cross-linking and thus effect on cuticle permeability remains to be better understood.

Nevertheless, while we targeted CDEF1, CDEF2 and CDEF3 for their putative involvement in the cuticle assembly, we concluded a function that is not common to other gene members of the related CDEF family. Indeed, in addition to the cutin hydrolytic activity associated with *At*CDEF1, several other CDEF1 close orthologs such as *Ca*GL1, *Ca*GLIP1, *At*GDSL1 and JNP1 have also shown lipase activity in different species, however described in a scope of defense to stress-related enzymes rather than cuticular assembly-related perspectives (sup. fig. 8) (Takahashi et al. 2010; Kim et al. 2008; Kram et al. 2008; Hong et al. 2008; Ding et al. 2020; Ding et al. 2019). Although conclusions of these studies indicate a role in preventing biotic or abiotic stress and enhancing plant resistance, investigations of cuticle of mutants would be interesting as it represents the first barrier to the environment. The other close ortholog *AtGGL22* has been associated to water retention in a stomatal-related study with mutants exhibiting an increase of stomatal density (Xiao et al. 2021). This result supports an important role in the organization of the plant surface which likely involves the remodeling of the cuticle such as the CDEF genes.

Interestingly, *At*CDEF1 did not show esterase activity toward tomato native cutin in our study, although it hydrolases the 2-MAG precursor and has been largely used for ectopic gene expression in roots to generate defect in suberin and cutin deposition (Berhin et al. 2019; Naseer et al. 2012; Li et al. 2017a; Ursache et al. 2018; Barberon et al. 2016; Wang et al. 2019). Differences in composition exist between the two species and their lipidic polymers, thus implying substrate specificities that corroborate research of Berhin et al. (2019). Although *CDEF1* has a different function, one cutin monomer was mainly affected in the mutant contrary to other cutin-related gene mutants, adding relevance to the question of substrate specificity that should be addressed in further studies.

### GDSL-hydrolase superfamily is largely involved in extracellular cuticle formation

The presence of *CUS* genes in genomes emerged during plant terrestrialization and likely appeared coterminous with a basic cuticle machinery to form the first plant cuticle (Jiao et al. 2020; Philippe et al. 2020; Kong et al. 2020). Multiplication of homologs has been described in seed plants, which lead to important family up to 14 genes in some species (Kong et al. 2020). Only two *CUS* genes are found in the early land plant *Physcomitrella patens* (bryophyte), but a double knockout is needed for the plant to exhibit a cuticle-related phenotype (Renault *et al*. congress communication). Comparably, the suberin assembly of *A.thaliana* requires the combinated knockout of at least 5 *SUS* genes to observe a phenotype (Ursache et al. 2021). Although the CUS family consists of five members in tomato, the fruit has been a particularly interesting model to study the function of these genes considering the massive expression of *CUS1* and the strong reduction of cutin deposition associated with its simple knockout (Girard et al. 2012; Yeats et al. 2012). Thus, the phenotype of *cus1* indicates that no homolog is able to compensate the important activity of CUS1 in the cutin assembly. Instead, members of the family have shown distinct expression profiles that reveal the relative importance of each of them in different development stages, tissues and organs. The cutin monomers composition varies depending on the tissue (*e.g.* dihydroxyhexadecanoic acid is major monomer in outer epidermis while hexadecenoic acid in inner epidermis), changing the physico-chemical nature of the assembly, which might affect the optimal activity of the CUS enzymes or suggests a substrate specificity. The different levels of polymerization activity among the CUS enzymes toward a single precursor substrate could not settle the question as both substrate specificity and the environmental conditions of the experiment might be involved. However, the unchanged relative proportions of cutin monomers in *cus* mutants (both outer and inner epidermis) as well as the highly conserved sequences of the CUS proteins imply that they are less subject to substrate specificity than physico-chemical environment of the cell wall and its degree of cutinization for their activity.

Beyond the CUS family, which is solely responsible for the polymerization of the cutin, we demonstrated that other GDSL-hydrolases also display this capability through *in vitro* assay. CDEF1, CDEF2 and CDEF3 can polymerize cutin precursors as a step of the remodeling activity and their family and 7 of the 9 members of the family may share this function as they have epidermis expression. Besides, tomato orthologs of *At*SUS enzymes form a neighbor family in the clade VI and are likely to share a lipid polymerization activity, although targeting different precursors than cutin (Yang et al. 2010). Considering that structural differences exists between all these families of enzymes and that slight differences might have significant effects on substrate specificity of GDSL-hydrolases (Wang et al. 2023), an important question for future studies is to determine the protein structural features that differentiate those closely related functions.

To date, no other GDSL-hydrolases have shown cutin polymerization activity, while hydrolysis of lipidic esters had been reported several times in the literature, and does not seem to be restricted to a specific clade. After *CUS1*, the 3 most expressed GDSL-hydrolase genes in tomato fruit epidermis are distant from the CUS/CDEF clade VI and remain to be characterized for their likely important roles considering their high expression. The fact that CUS enzymes belong to the GDSL-hydrolases that catalyzes various reactions on different substrates makes this large superfamily interesting to decipher mechanisms related to the cuticle assembly that remain largely unknown. Catalyzing speculative linkages of cutin with phenolic or polysaccharidic compounds are among putative function to explore, as well as involvement in the rearrangement of cutin and polysaccharides in cuticle during developmental growth (Reynoud et al. 2022). Considering that GDSL-hydrolase orthologs often retain acquired or lost function following evolutive event (Volokita et al. 2011), the highly expressed SFAR4 would be an interesting candidate for a such role as substrates related to phenylpropanoid metabolism and cell-wall are identified for enzymes located in GDSL-hydrolase clade IV. Several interesting candidates from the clade V also raise particular interest for putative roles in cuticle formation, including the OPS1 that is a stomata-related ortholog highly expressed in a stomata free model, and should also be considered for investigation in further research.

## Conclusion

Number of GDSL-hydrolases are specifically expressed in plant epidermis, while a wide range of functions are associated with enzymatic activities in this superfamily of enzymes. In this study, we investigated the role of candidate genes in the cuticle assembly, for which mechanisms of extracellular formation remain largely unknown. We reported a spatio-temporal coordination of GDSL-lipases enzymes in the context of fruit growth, including cutin biosynthesis and remodeling, the latter function being assigned for the first time to an identified enzyme. In addition, our study is the first investigating of region-specific gene expression and cutin polymerization at the surface of the fruit, which represents a step toward the understanding of the cuticle assembly integrating the biology of the fruit.

## Acknowledgements

We are grateful to Prof. Maria Alejandra Gandolfo Nixon and Jennifer L. Svitko for technical assistance and access to the SEM, as well as Anthony M. Condo for technical assistance to the MALDI-TOF. We thank Lily E. Duplooy, Milay M. Haskin, Annelise Viera and Stephen Snyder for their assistance and help with time consuming experimentations.

## Author Contributions

GP, IS, and JKC designed the research. GP, IS, AG, MJC, DSD and MHC performed experiments and analyzed the data. GP, IS and JKC and wrote the paper.

## Table legend

Ranking of the expressed *GDSL-hydrolase* genes in tomato fruit epidermis (>1rpm value in outer or inner epidermis from the TEA database). Bold genes are selected for recombinant protein production in this study; asterisks indicate specific epidermis expression genes; Other bold data indicate spatiotemporal interest of CDEF and CUS genes; localization of highest gene expression on fruit surface is taken from total pericarp data at mature stages (data unavailable for growing stages); downregulated genes in *gpat6* tomato mutant are indicated from Petit et al. (2016).

