## Supplemental data for "Spatiotemporal variation in cutin polymerization and remodeling mediated by GDSL-hydrolase enzymes during tomato fruit development"

#### Supplemental Figure Legends

Supplemental figure 1- Heatmaps of the relative expression of selected candidate GDSL-hydrolases in different fruit tissues (vertical axis) and developmental stages (horizontal axis), based on the TEA database (https://tea.solgenomics.net/). The color gradient indicates the relative gene expression levels from white (lowest) to brown (highest), based on RNA-seq analysis of the distinct pericarp cell types.

Supplemental figure 2- Enzymatic incubation of 2-MAG with recombinant purified enzymes. (A-B) 12h reaction of successive incubation of CUS1 and a candidate GDSL-hydrolase. Deactivation step of CUS1 by boiling the mixture occurred at different time before re-incubation (around 4h in A and 6h in B). (C) Incubations of CDEF enzymes with 2-MAG for 24h. (D) Glycerol release assays perform on enzymatic mixtures after 6h incubations.

Supplemental figure 3- Semi-*in vivo* bioassay screening of cutin polymerization activity with purified enzymes. Inner epidermis of cut-in-half 15dpa tomato fruit M82 (WT) are incubated with purified recombinant GDSL-hydrolases (red dots) or commercially pure *Fusarium solani* cutinase (blue dots) for 12h, then stained with Toluidine Blue.

Supplemental figure 4- Genome edited sequences by CRISPR/Cas9 on M82 tomato genotype. GDSL motif surrounded by orange frames, CRISPR targets highlighted in green and blue for *CDEF* genes and *CUS2* respectively ; deletion or addition are restricted to exon sequences and colored in red, leading each time to shifting of translation frame and gene knockout.

Supplemental figure 5- Screening for fruit inner epidermis phenotype of red ripe stage for *cus2* genotypes. (A) Toluidine Blue staining of inner epidermis does not indicate higher cutin permeability in *cus2* lines than their respective genomic background (M82 or *cus1*). (B) Monomer profile of inner epidermis cutin of *cus2* lines is not significantly different than their respective genomic background. *, $P < 0.05$; **, $P < 0.01$ (Student's t test).

Supplemental figure 6- Screening for fruit cuticle phenotype in stem, equatorial and stylar regions of red ripe stage fruit for *cdef1/2* genotypes. (A) Cutin monomer composition of isolated cuticle. (B) Cuticle thickness measurement of epidermis cross-section. (C) Water permeance of isolated pieces of cuticles isolated with no significant differences observed between WT and mutants lines. Statistical differences are indicated by letters given by Tukey's HSD tests following significant ANOVA results.

Supplemental figure 7- Measurement of Young’s modulus (A) and maximum force at breaking point (B) on isolated and dewaxed cuticle samples. No significant differences have been found through ANOVA test.

Supplemental figure 8- Maximum Likelihood phylogenetic tree of GDSL-hydrolases from the clade I including the studied CUS1 and CDEF enzymes from tomato (names in blue) and a selection of GDSL-hydrolases from other species (names in green). Bootstrap values above 70 are indicated on tree branches.

### Supplemental figure 1

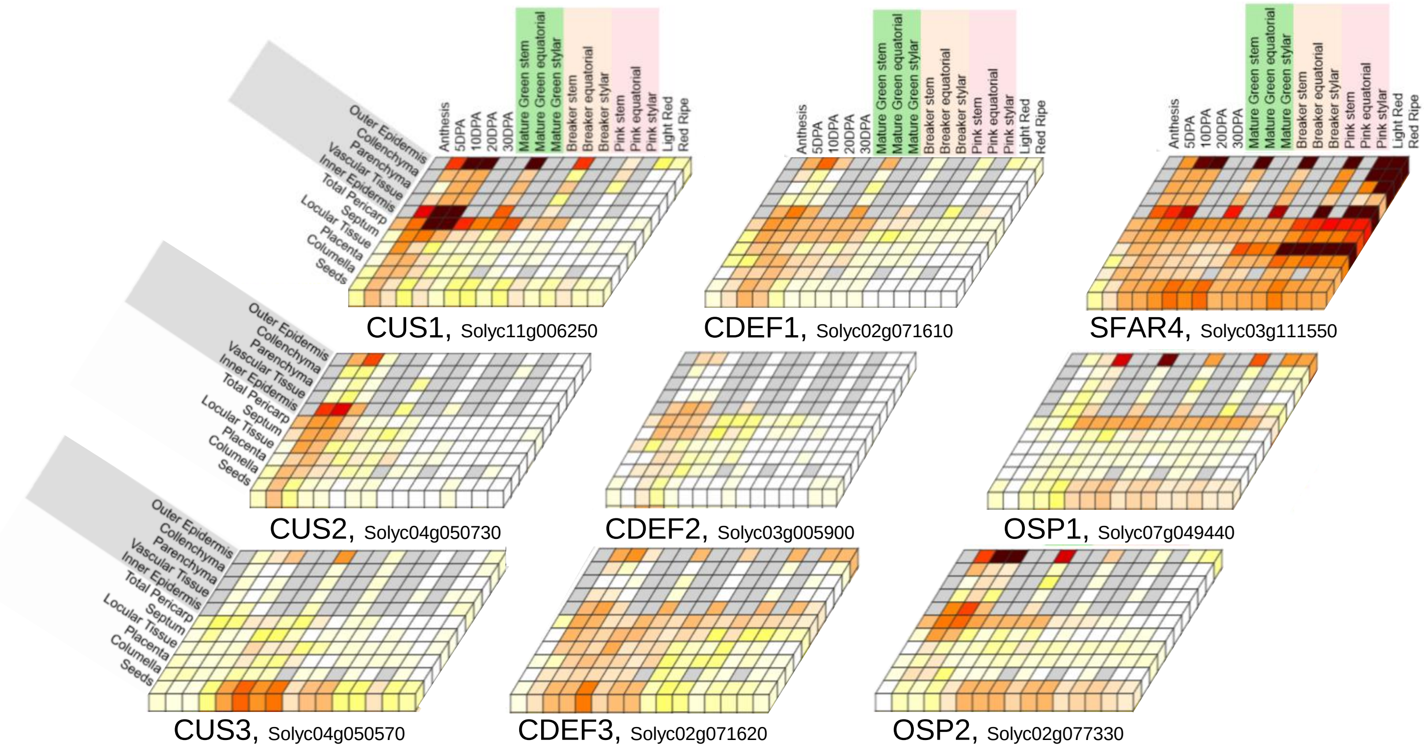

### Supplemental figure 2

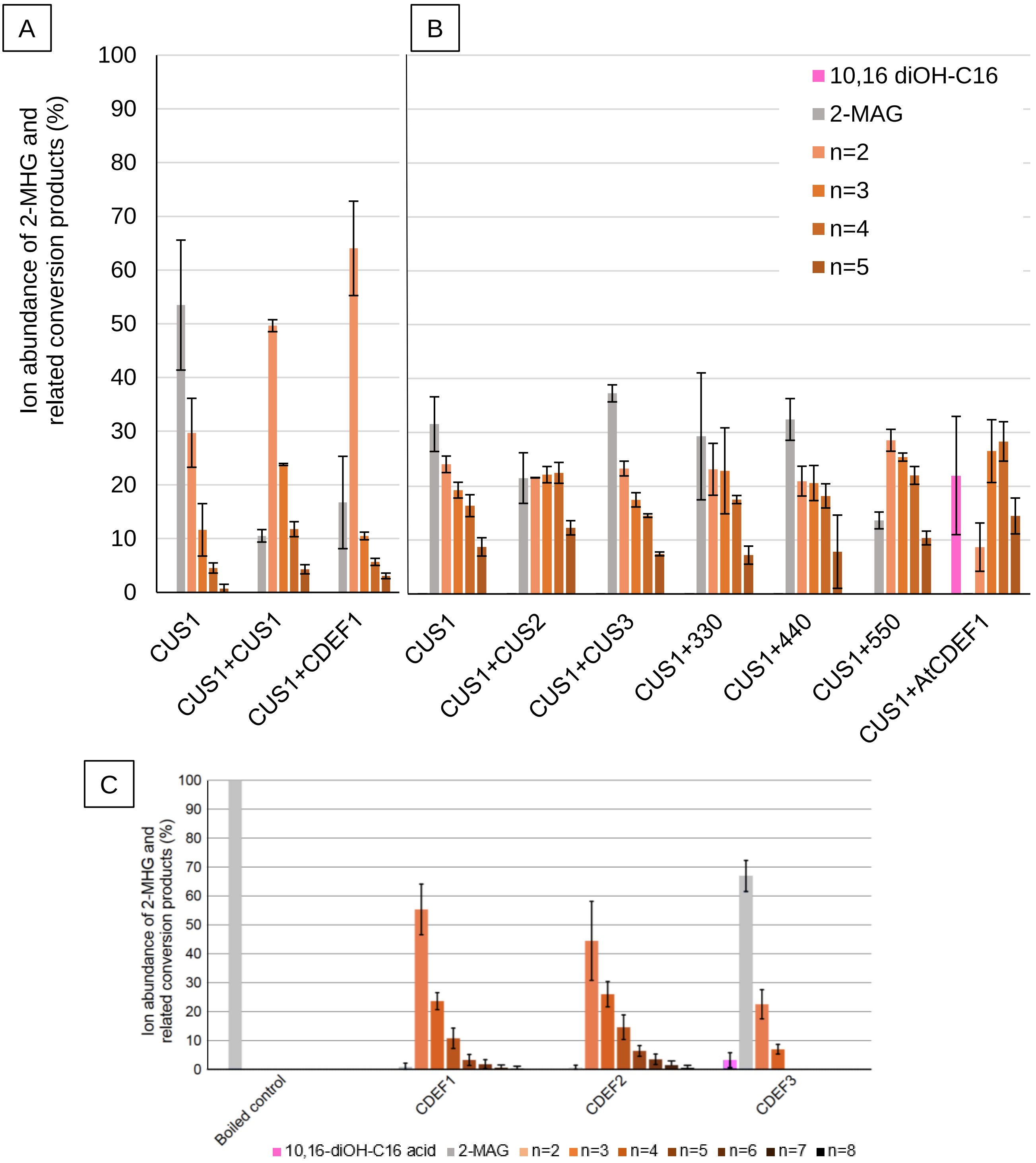


D

| | CUS1 | CUS2 | CDEF1 | *At* CDEF1 |
|---|---|---|---|---|
| nmol/well | 6,82 | 1,83 | 20,85 | 18,69 |

Supplemental figure 3

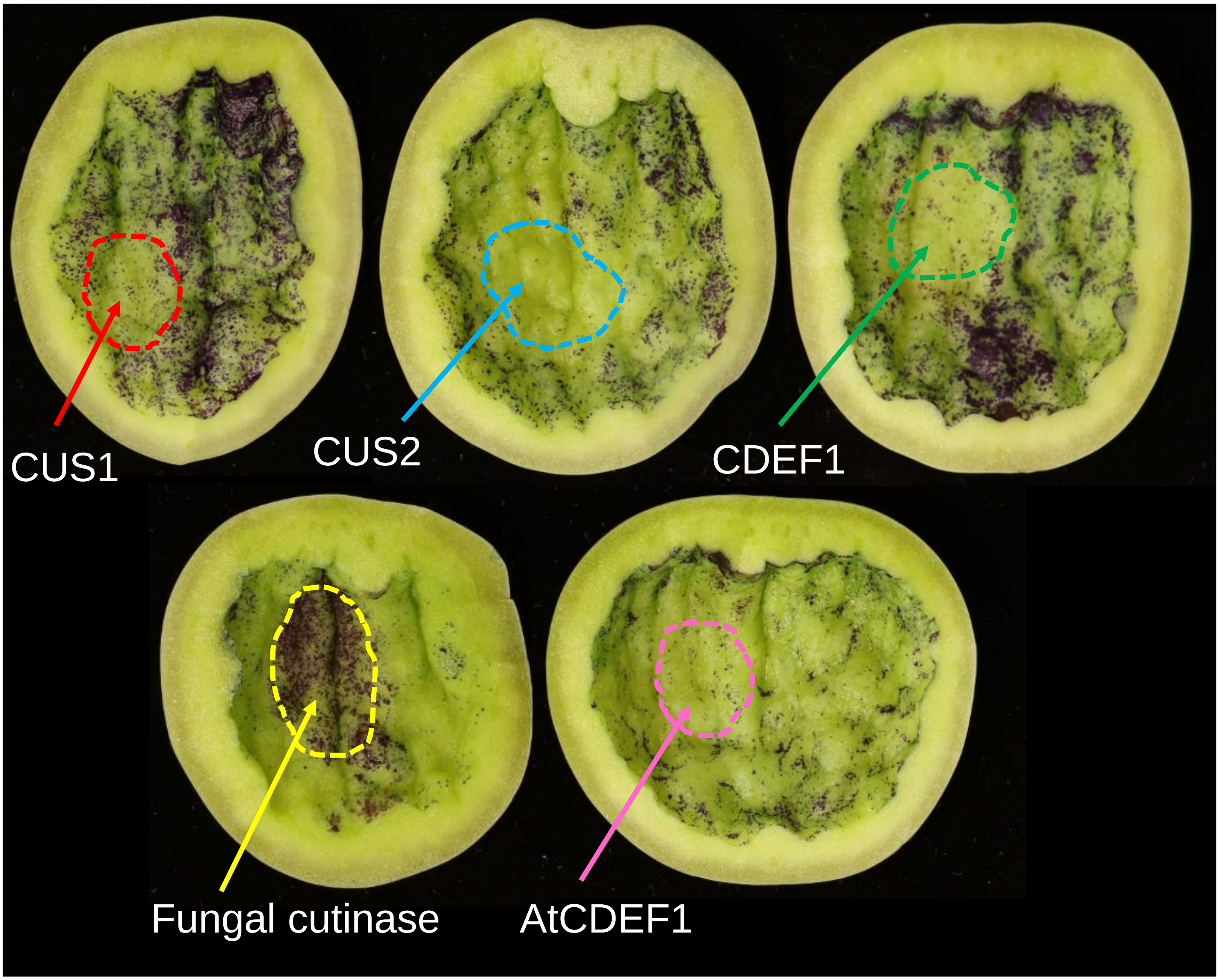

### Supplemental figure 4

- *SlCDEF1* mutations

**M82** TTGGTGATTCATTAGTGGATAATGGGAATAACAACAATATTCAGTCATTGGCTAGGGCTAATTACTTGCCTTATGGTATTGATTATCCTGATGGTCC-AACTGGAAGGTTTTCCAATGGAAAAACAACTGTTGATGTCATTG
***cdef1/2*-L1** TTGGTGATTCATTAGT---TAATGGGAATAACAACAATATTCAGTCATTGGCTAGGGCTAATTACTTGCCTTATGGTATTGATTATCCTGATGGTCCCAACTGGAAGGTTTTCCAATGGAAAAACAACTGTTGATGTCATTG
***cdef1/2*-L2** TTGGTGATTCA--------------------------------------------------------------------------------------------------------------------------------TTG

- *SlCDEF2* mutations

**M82** AGGAGAACCACAAGTGCCTTGTTACTTTATATTTGGTGATTCATTAGTGGATAATGGAAATAATAATGTGATTAGGTCATTGGCTAGAGCTGATTATTTGCCTTATGGAATTGATTTTCCAGATGGACCAACTGGAAGATT
***cdef1/2*-L1** AGGAGAACCACAAGTGCCTTGTTACTTTATATTTGGTGATTCAT-------TAATGGAAATAATAATGTGATTAGGTCATTGGCTAGAGCTGATTATTTGCCTTATGGAATTGATTTTCCAGATGGACCAACTGGAAGATT
***cdef1/2*-L2** AGGAGAACCACAAGTGCCTTGTTACTTTATATTT-----------------TAATGGAAATAATAATGTGATTAGGTCATTGGCTAGAGCTGATTATTTGCCTTATGGAATTGATTTTCCAGATGGACCAACTGGAAGATT

- *SlCUS2* mutations

**M82** AAGGTCGAGCCTTCTTCGT-GTTTGGTGATTCACTAGTAGATAACGGGAACAATAATTACTTAGTTACTAGTGCCAGGGCAGATTCTCCCCCCTATGGCATTGACTATCCTACTCATCGTGCCACAGGTCGATTTTCCAATGGACT
***cus2*-L1** AAGGTCGAGCCTTCTTCGT-GTTTGGTGATTCACTAGTAGATAACGGGAACAATAATTACTTAGTTACTAGTGCCAGGGCAGATTCTCCCCCCTATGGCATTGACTATCCTACTCATCGTGCCACAGGTCGAT----CAATGGACT
***cus2*-L2** AAGGTCGAGCCTTCTTCGTTGTTTGGTGATTCACTAGTAGATAACGGGAACAATAATTACTTAGTTACTAGTGCCAGGGCAGATTCTCCCCCCTATGGCATTGACTATCCTACTCATCGTGCCACAGGTCGATTTTCCAATGGACT

### Supplemental figure 5

A

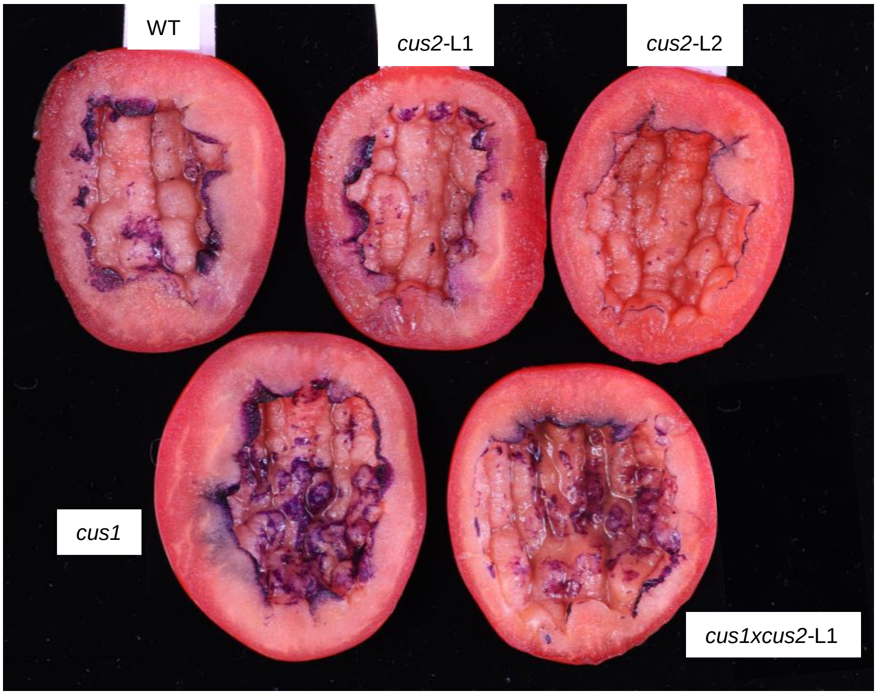


B

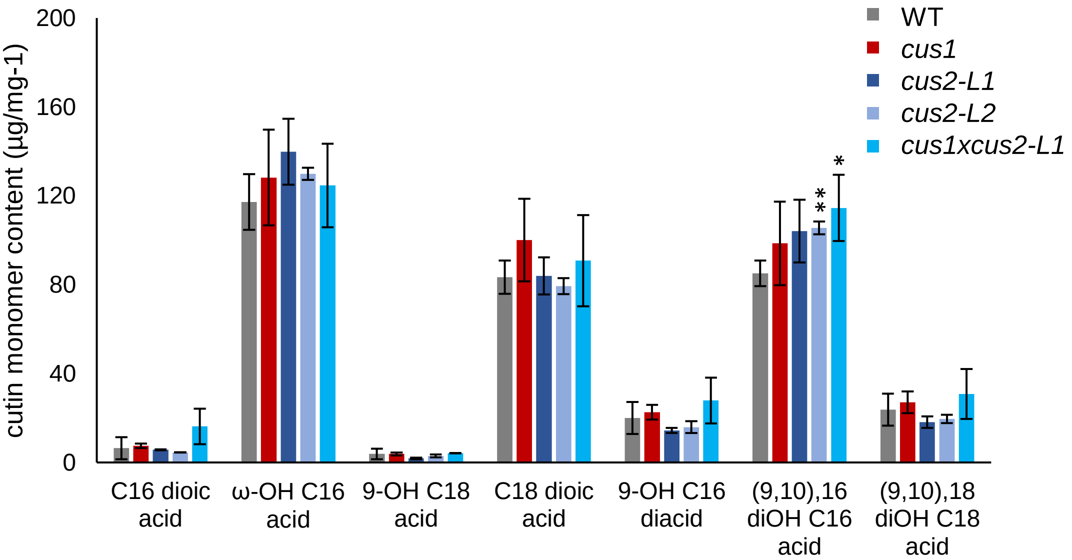

### Supplemental figure 6

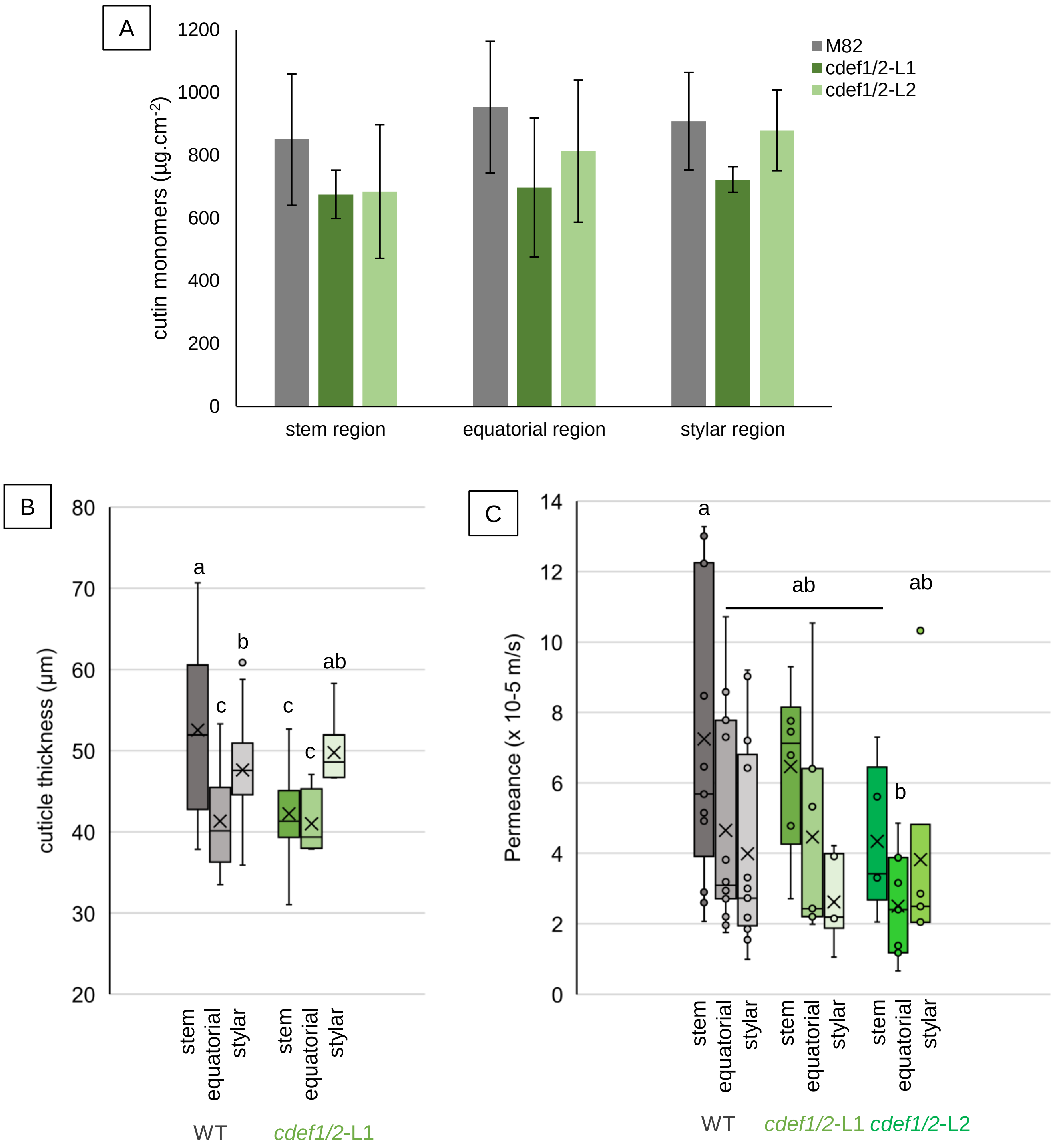

### Supplemental figure 7

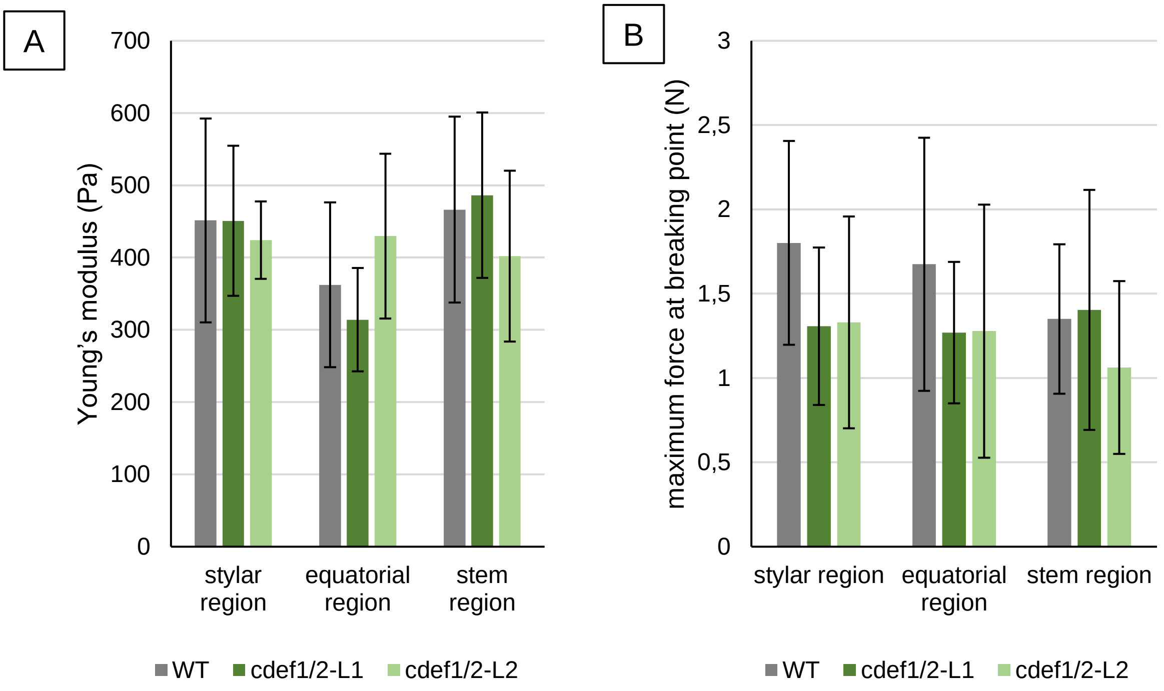

Supplemental figure 8

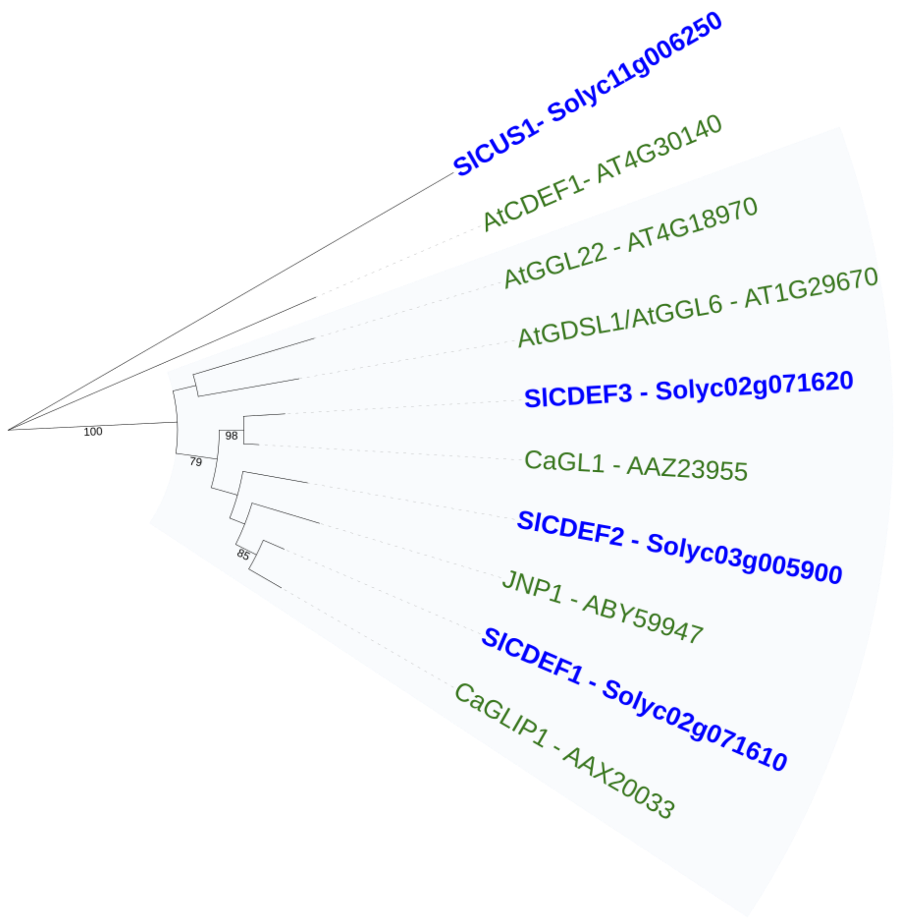
